# Defining the Yeast Resistome through *in vitro* Evolution and Whole Genome Sequencing

**DOI:** 10.1101/2021.02.17.430112

**Authors:** Sabine Ottilie, Madeline R. Luth, Erich Hellemann, Gregory M. Goldgof, Eddy Vigil, Prianka Kumar, Andrea L. Cheung, Miranda Song, Karla P. Godinez-Macias, Krypton Carolino, Jennifer Yang, Gisel Lopez, Matthew Abraham, Maureen Tarsio, Emmanuelle LeBlanc, Luke Whitesell, Jake Schenken, Felicia Gunawan, Reysha Patel, Joshua Smith, Melissa S. Love, Roy M. Williams, Case W. McNamara, William H. Gerwick, Trey Ideker, Yo Suzuki, Dyann F. Wirth, Amanda K. Lukens, Patricia M. Kane, Leah E. Cowen, Jacob D. Durrant, Elizabeth A. Winzeler

## Abstract

*In vitro* evolution and whole genome analysis were used to comprehensively identify the genetic determinants of chemical resistance in the model microbe, *Saccharomyces cerevisiae*. Analysis of 355 curated, laboratory-evolved clones, resistant to 80 different compounds, demonstrates differences in the types of mutations that are identified in selected versus neutral evolution and reveals numerous new, compound-target interactions. Through enrichment analysis we further identify a set of 137 genes strongly associated with or conferring drug resistance as indicated by CRISPR-Cas9 engineering. The set of 25 most frequently mutated genes was enriched for transcription factors and for almost 25 percent of the compounds, resistance was mediated by one of 100 independently derived, gain-of-function, single nucleotide variants found in 170-amino-acid domains in two Zn_2_C_6_ transcription factors, *YRR1* and *YRM1* (p < 1x 10 ^−100^). This remarkable enrichment for transcription factors as drug resistance genes may explain why it is challenging to develop effective antifungal killing agents and highlights their important role in evolution.

## Introduction

Over the past decade, decreases in sequencing costs have led to an explosion in the number of cataloged genetic variants in all fields of biology. In a recent sequencing study of 1100 yeast isolates 1,625,809 single nucleotide variants (SNVs) were identified^1^. Sequencing of thousands of mosquitoes that cause human malaria identified 57 million variants^2^. There are now 660 million cataloged human variants^3^. The challenge lies in efficiently identifying variants that change the phenotype of tumor, pathogen or agricultural pest especially when the genetic background is heterogenous and there may be tens of thousands of differences between two sequenced isolates.

Systems for studying what these SNVs do for the cell have lagged. Systematic functional genomic studies have continued to rely on strain libraries in which the entire coding region is modified. For example, after the yeast genome was sequenced a set of homozygous and heterozygous knockout strains was constructed which bear deletions in all genes in the genome^4,5^. This set has been used to repeatedly and systematically identify knockout/knockdown lines that show sensitivity or resistance to a wide variety of different compounds^6^ and remains important^7^. CRISPR-based, genome-wide knockout and knockdown studies are now being employed in many organisms to identify drug targets^8^ or study processes such as the emergence of cancer drug resistance. Such studies will miss gain-of-function SNVs, which often drive natural adaptive evolution.

Although systematic CRISPR-Cas9 based analyses of SNVs are also feasible^9^ with the reduction in costs of whole genome sequencing, experimental evolution, which mimics natural evolution, becomes more attractive. Here we exposed the model yeast to a large set of compounds, similar to cell-permeable small molecules or natural products that fungi might encounter in their natural environment or which might be used in agriculture or medicine. Whole genome sequencing of 355 curated, evolved, compound-resistant clones showed only a few new coding variants per clone and that statistical approaches can be used to readily identify variants that modify phenotype. We discover an enrichment for gain-of-function variants that affect transcription. These data may provide clues about why it is challenging to develop small molecule therapeutics against fungi.

## Results

### Building a library of compounds that are active against a drug-sensitive yeast

To understand how yeast evolve to evade the action of small molecules, we first assembled a collection of molecules and evaluated their activity against the yeast *S. cerevisiae*. Specifically, we tested compound libraries comprised of (1) drugs approved for human use, (2) well known tool compounds, and (3) compounds from open-source libraries with demonstrated activity against other eukaryotic pathogens, viruses, or tuberculosis. Commercially available compounds were tested in dose response, while members of other libraries were initially tested at a single point concentration of 150 μM in biological duplicates. Those that showed at least 70% growth inhibition were subsequently tested in dose response. To increase the probability of finding compounds with activity against yeast at physiologically-relevant concentrations, and given that compound cost, availability and resupply are major impediments to a study such as this, we used a sensitized strain of yeast, termed the “green monster (GM)” in which a variety of ABC transporters have been replaced with GFP^10^.

Overall, the compounds of the assembled collection had drug-like physiochemical properties in terms of molecular weight and the number of hydrogen bond donors and acceptors (**Figure 1A**). Maximum Common Substructure (MCS) clustering identified 307 clusters with a similarity coefficient of 0.64 (**Figure 1B, Table S1**). Altogether, 286 compounds had an IC_50_ < 76 μM, and 98 compounds had an IC_50_ < 10 μM (**Table S1**). Of these 286 active compounds, 165 share a MCS with at least one other compound (**Table S1**). Cluster enrichment was observed at rates greater than expected by chance for some compounds. For example, there were 12 members of a benzothiazepine family (cluster 185, **Figure 1B**) in the tested group of 1,618 compounds of which six had an IC_50_ of less than 10 μM in the yeast model (p = 2.46 × 10^−5^).

**Figure 1:**
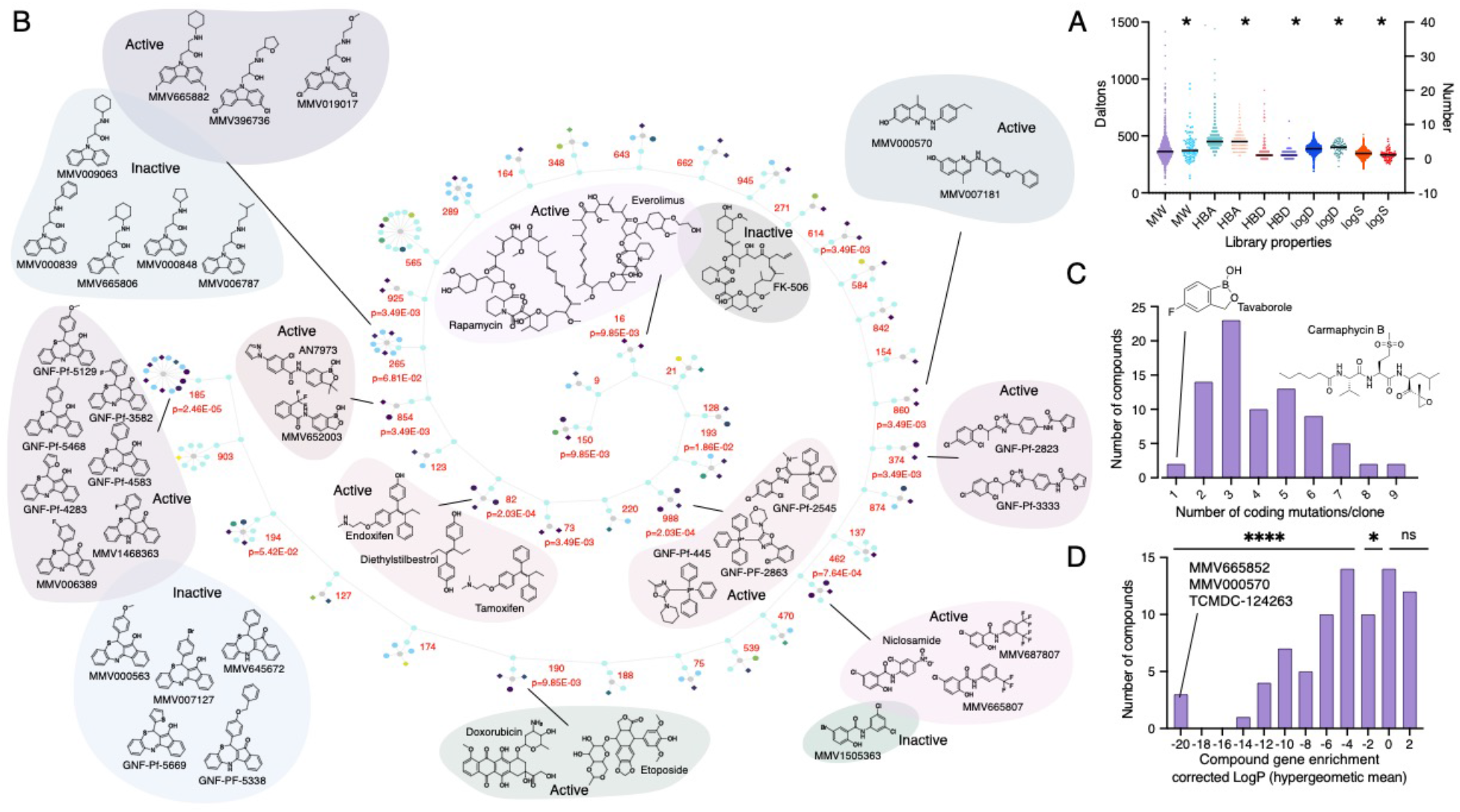
Compound Summary. **A.** Lipinski properties of compounds used in this study calculated calculated using StarDrop version 6.6.4. Left Y axis: MW, molecular weight: Right Y axis: HBD, hydrogen bond donor, HBA, hydrogen bond acceptor, logD, logS. * indicates 80 compounds that yielded resistant clones. **B. Maximum Common Substructure clustering analysis for 80 compounds yielding resistant clones and larger library of 1600**. The diagram shows 41 clusters sharing a Maximum Common Substructure (MCS) from which at least one compound was selected for drug response (indicated by diamonds). Circles represents compounds that were not selected or were inactive. The strength of cytotoxicity against the *S. cerevisiae* GM strain of tested compounds is indicated by the node’s color intensity from purple (higher potency) to yellow (lower potency). Probability values were calculated using the hypergeometric mean function showing that enrichment for clusters was greater than expected by chance. Compounds from clusters with a p-value of less than 0.05 and which had multiple members active against GM are shown. **C. Coding region mutations for selected compounds.** Histogram showing the distribution of the number of coding mutations (e.g. missense, start-lost) per clone for the set of 80 compounds used in selections. **D. Compound gene enrichment for selected compounds.** Histogram showing the distribution of the probabilities of discovering multiple hits in a single gene per compound, calculated using a Bonferonni-corrected hypergeometric mean function for the set of 80.

### In vitro resistance evolution and whole genome analysis link compound structures to phenotype

Based on potency and compound availability, we selected 100 active compounds from this collection for follow-up *in vitro* evolution experiments. To select for resistant strains, yeast cultures (approximately 10^8^-10^9^ cells) were first inoculated with sublethal compound concentrations. We subsequently ramped up the selection pressure by serial dilution of the saturated cultures into media with increased compound concentrations until resistance was observed as measured by an increase in IC_50_ values. Using this strategy, we successfully isolated 355 strains resistant to 80 compounds. Cultures were considered resistant if they (1) continued to grow at compound concentrations above the IC_50_ value of the untreated culture, and (2) had at least a 1.5-fold shift in IC_50_ value compared to the drug-naïve parental line (**Table S2**). These resistant cultures were plated on drug-containing plates to isolate single colonies. Genomic DNA was isolated from these clones, and whole genome sequencing of both the resistant and parental strains identified mutations associated with resistance. The IC_50_ values of the resistant clones increased 1.5- to 5-fold for 121 resistant strains, 5- to 10-fold for 101 resistant clones, and > 10-fold for 98 resistant strains (**Table S2**). In about 20% of the cases, we were unable to isolate resistant strains after many weeks of selections, often because of contamination or poor compound availability.

Next clones were sequenced to 55-fold average coverage (**Table S3**) using short read methodology. To detect mutations, we designed a custom whole genome analysis pipeline and filtering method (see Methods) that was automated through the computational platform, Omics Pipe^11^. Briefly, raw sequence reads were aligned to the *S. cerevisiae* 288C reference genome (assembly R64). SNVs and INDELs were called using GATK HaplotypeCaller^12^, filtered to retain only those of high quality and high allele fraction (appropriate for a haploid organism) and annotated with SnpEff^13^. Mutations were only considered to be potentially resistance-conferring if they were present in the evolved clone but not in the drug-sensitive parent. We discovered 1,405 high quality mutations (1,286 SNVs and 119 indels) in 731 unique genes that arose during the course of drug selection, with an average of 3.96 mutations per line (**Table S4).** On average we observed between 1 to 8 coding mutations per evolved clone per compound (**Figure 1C**) with some variation. For example, selections with the small synthetic antifungal, tavaborole, produced four resistant clones (8- to >15-fold increase in IC_50_) with six total missense mutations (four in the target), while selections with the natural product, carmaphycin, resulted in three resistant clones with 28 mutations, of which 22 were coding (one in the predicted target). For the majority of compounds, we observed a strong enrichment and reproducibility for specific genes (**Figure 1D**). For example, for compound MMV665852 we obtained 10 independent resistant clones with 29 independent coding mutations, of which seven were in a single gene, *YRR1*. Given that yeast has roughly 6,000 genes, the Bonferonni-corrected probability of this enrichment by chance is roughly 8.38 × 10^−21^.

To further assess the likelihood that our evolved mutations would contribute to resistance we considered the types of mutations and compared these to a published set of 3,137 neutral mutations in yeast strains grown long term without compound selection^14^. We observed significantly different distributions in our drug-selected set than in the neutral set (χ^2^, *p* < 0.0001). For example, 39% of the nucleotide base transitions for selected 1,286 single nucleotide variants (SNVs) were C to A or G to T, while for the 3,137 neutral transitions, 40% were for A to G or T to C (**Figure 2A**). These data provide additional evidence that the observed mutations provide a selective advantage to the evolved clones. Likewise, we observed a noteworthy difference in the coding changes. Among exonic selected mutations, 994 were nonsynonymous and 127 were synonymous (~8:1 ratio), indicating that drug treatment applied a strong positive selection, as expected (**Figure 2B**). In contrast, the neutral SNVs had a strong bias toward synonymous mutations (**Figure 2C**).

**Figure 2:**
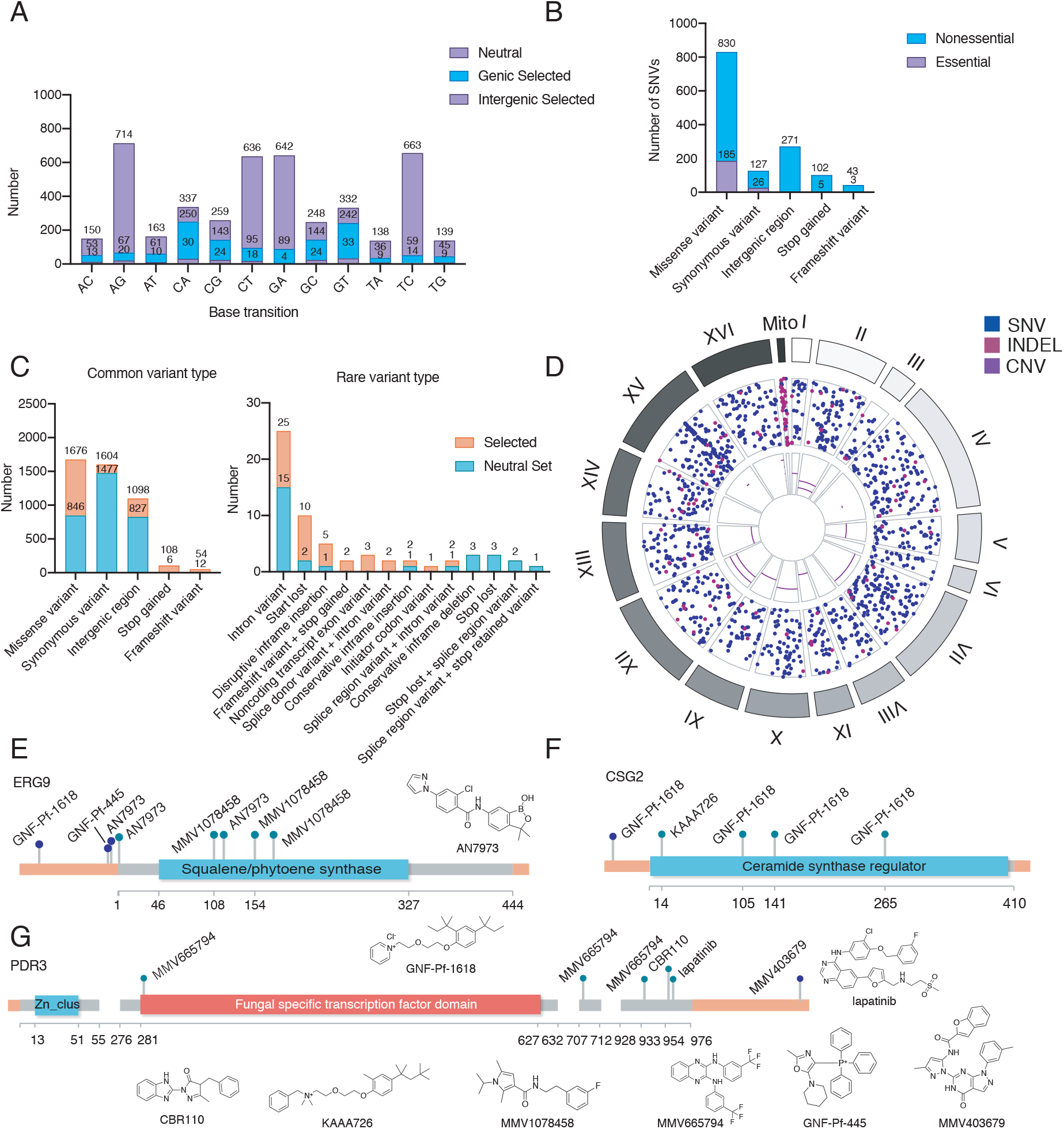
Mutations observed in yeast IVIEWGA experiments. **A**. **Classification of mutation based on base transition type for 1286 SNV mutations obtained via compound selection and neutral mutations ^14^. B**. **Classification of mutation types and their occurrence in essential vs. nonessential genes.** Essentiality data was imported from the *Saccharomyces* Genome Deletion Project database. **C. Mutation type.** Variant classes for 1405 mutations (indels and SNVs, Supplemental Table S4) obtained via compound selection versus neutral mutations ^**14**^. **D**. **Circos plot**. Circos plot of SNPs (blue), INDELs (magenta), and CNVs (purple) identified through resistance generation, generated with BioCircos R package^73^. **E-G. Intergenic mutations**. Plot locating each coding region mutation onto the gene (grey) and intergenic mutation onto the chromosome (orange) based on the calculated distance.

Copy number variants (CNVs) were also detected through a coverage-based algorithm using the output from GATK DiagnoseTargets. We observed 24 CNVs in our resistant strains (**Table S5**). Unlike for studies in *P. falciparum*, we frequently observed aneuploidy (11 times) in addition to small, discrete, intrachromosomal amplifications (13 times). Altogether, we observed aneuploidy with eight compounds, including BMS-983970, doxorubicin, etoposide, GNF-Pf-3582, GNF-Pf-4739, hygromycin B, CBR110, and wortmannin (**Figure 2D**). This is perhaps not surprising given that aneuploidies arise in *S. cerevisiae* as a short-term stress response^15^. We observed an amplification on chromosome XVI that involved the bZIP transcription factor, *ARR1,* for clones resistant to GNF-Pf-1618 and GNF-Pf-2740, as well as with four strains resistant to MMV665794. The strains, Wortmannin-13R3a and sBMH113-7R4a, both had chromosome XV CNVs that involved the transcription factors *YRR1* and *YRM1* (discussed below).

Our large mutational dataset also offers broad insights into the functional impacts of different variant types. Synonymous and missense variants emerged in essential genes in approximately 20% of cases. This finding agrees with the literature, which suggests that only 20% of the yeast genome encodes essential genes^16^. By this same metric, mutation types with more disruptive impacts on the resulting protein, such as premature stop codons and frameshift variants, deviate strikingly from the expected genome-wide value of 20%. These mutations occur in essential genes only 4.9% and 7.0% of the time, respectively (**Figure 2B**). Of these ten disruptive mutations four were in a single gene, *PAN1*, involved in the regulation of endosome internalization.

### Resistance-conferring intergenic mutations are rare

Although intergenic mutations are frequently found in cells not subject to selection, mutations in promoters or 3’ UTRs might confer resistance by increasing or decreasing transcript levels. We mapped intergenic mutations to their nearest-neighbor coding genes (**Table S6**). In contrast to coding mutations, where mutations in specific genes appeared repeatedly, this analysis showed little enrichment. However, we did observe several repeated mutations in the intergenic promoter regions of a few genes. For example, we discovered three mutations upstream of the ergosterol biosynthesis and azole resistance gene, *ERG9*^17,18^, in addition to seven *ERG9* coding region mutations. One of the intergenic mutations of *ERG9* falls in the putative promoter region and was observed in selections with compound AN7973, which was also associated with two mutations in the coding region (**Figure 2E**). Four coding mutations and one non-coding mutation upstream of the starting codon in the endoplasmic reticulum membrane protein and capsofungin resistance protein^19^, *CSG2,* were also observed. All five mutations (including the intergenic mutation) are associated with selections to compound GNF-Pf-1618 and its close analog, KAAA725 (**Figure 2F**). We also identified five mutations in the coding region of *PDR3*^20^, a transcriptional regulator of the multidrug efflux, and an additional mutation was found downstream of the open reading frame (**Figure 2G**). These data show that most mutations identified in resistant clones are coding, although intergenic mutations should not be entirely dismissed.

### CRISPR/Cas9 validation shows that most genes identified more than once confer resistance, but singletons mutations may not

The presence of one or multiple SNVs in a resistant line is not proof that a specific mutation confers resistance since many mutations, so called hitchhiker mutations, can co-occur in a resistance strain and can even be non-adapative^21^. To confirm the role of these mutations in a clean genetic background, we used CRISPR/Cas9 technology to introduce 65 altered alleles from the evolved mutants back into the original (unevolved) strain. Successfully reverse-engineered strains were tested in liquid-growth assays using the same compounds from the corresponding IVIEWGA experiments. We compared the IC_50_ values of these strains to those of the parental strain using a cutoff of 1.6-fold shift between edited and parental line. In total, this comparison verified that 50 genetic changes representing 39 unique genes contributed to the observed resistance (**Table S7**). Mutations that were repeatedly identified tended to have a high probability of confirmation. The only exception was *RPO21*, a subunit of RNA polymerase, which was unconfirmed and mutated four separate times (two nonsynonymous and two synonymous mutations). For the 20 alleles that did not show a 1.6-fold change (**Table S8**), we noted that 11 of the resistant clones also carried additional resistance alleles in a highly represented gene such as *YRR1* or *YRM1* (**Table S4**). In addition, some of the “unconfirmed” CRISPR-Cas9 alleles resulted in a statistically-significant, gain of sensitivity. For example, a clone with an edited G454S mutation in *UTP18* is 2-fold and 4-fold more sensitive to MMV665852 and CBR868, respectively, than the parent, possibly because the resistant strain has slower growth but better survival in the presence of a cytotoxic compound due to another resistance mutation. In fact, it is known that mutations in *RPO21* result in transcriptional slippage, which may allow cells to better survive cytotoxic drugs that alter nucleotide pools^22^.

### Using in vitro evolution for drug target and mechanism of action studies

A major advantage of using yeast is that it is a model system for target discovery. To assess the functional importance of mutations we considered individual compounds and their mechanism of action. For compounds with defined targets we frequently identified mutations clustering in the active site of the proposed target molecule. For example, we isolated six strains resistant to flucytosine (**Table 1**). Of the nine identified missense or stop mutations, six were in the uracil phosphoribosyltransferase domain of *FUR1* (probability of enrichment by chance = 1.2 × 10^−26^ using hypergeometric mean function). A homology model (**Supplemental Figure 3A**) reveals that they are all located near the 5-FUMP binding pocket, suggesting that these changes confer resistance by disrupting 5-FUMP binding. We obtained four tavaborole-resistant strains that were highly resistant (> 15 μM). These strains only six new high allele-fraction SNVs, four of which (R316T, V400F, V400D, and M493R), all of which were coding mutations in the 145 amino acid aminoacyl-tRNA synthetase editing domain of *CDC60* (p = 1.11 × 10^−18^ hypergeometric mean function), the gene that encodes leucyl tRNA-synthetase in yeast. A LeuRS homology model (Supplemental **Figure 3B**) with a tavaborole ligand docked using QuickVina2^23^ suggests that the *CDC60* mutations confer resistance by directly interfering with tavaborole binding to Cdc60.

**Table 1.**
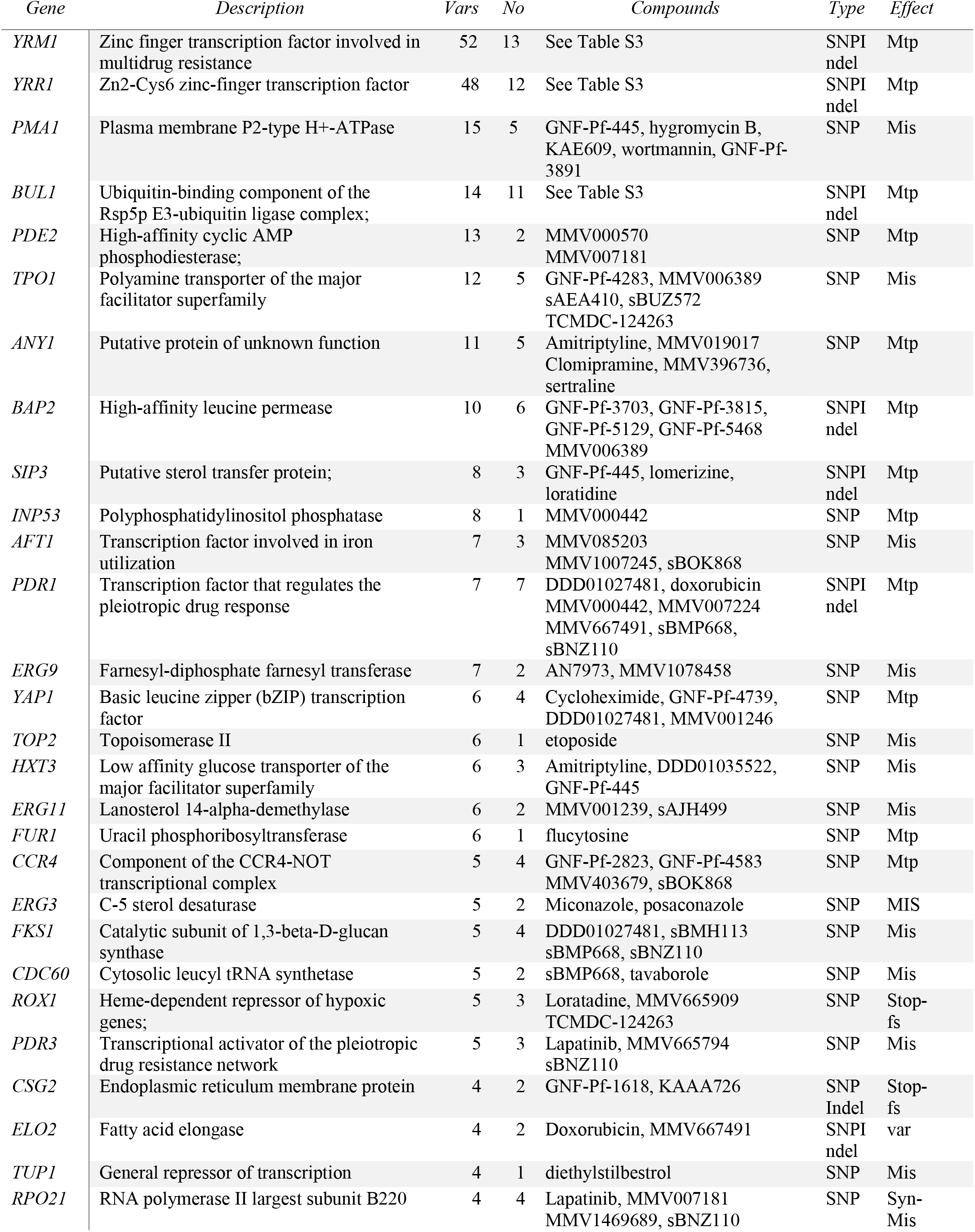
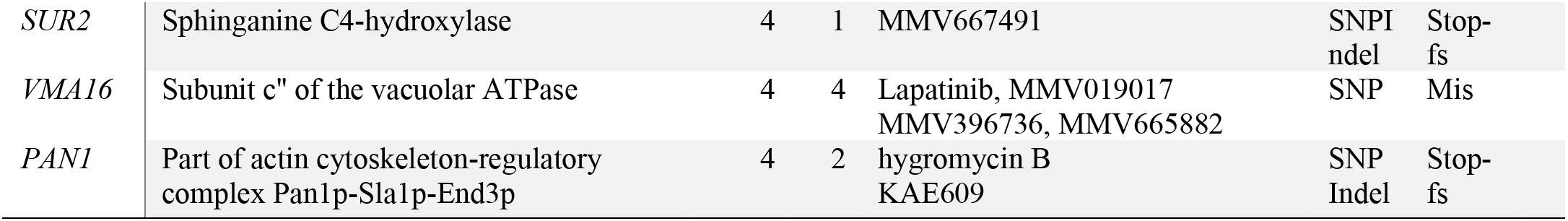
Genes detected at least 4 times *in vitro* evolution experiments. Vars = number of times gene was identified as mutated in independent evolution experiments. No. = number of compounds. Effect (fs, frameshift; mis, missense; syn, synonymous; mtp, multiple effects).

We also examine compounds that are used in chemotherapy. Camptothecin is a specific topoisomerase (Top1) inhibitor that binds the DNA/Top1 cleavage complex, preventing DNA religation^24^. We isolated two camptothecin-resistant yeast clones with three missense mutations, two of which were in *TOP1* (G297C and E669*) (**Table 1**). A homology model (**Figure 3C**)^25^ was constructed by aligning a partial yeast Top1 crystal structure to a crystal structure of the human TOP1 with camptothecin bound (PDB: 1T8I)^26^. This model showed that G297 is located in the core domain of the enzyme near the binding pocket, suggesting that it confers drug resistance by directly impeding drug binding, while E669* truncates the entire C-terminal domain, which contains the DNA-binding site^27^ (**Figure 3C**), thus eliminating many protein/DNA contacts and likely impeding the formation of the drug-DNA-protein complex. Rapamycin, a macrocyclic lactone, and its analog everolimus (a so-called rapalog), potently inhibit mTOR, a protein kinase component of both the mTORC1 and mTORC2 complexes that controls cell growth and proliferation in many species. Two rapamycin resistant and three everolimus-resistant clones were identified. One carried a (S1975I) mutation in the FKBP12-rapamycin binding domain of mTOR (**Table S4**) and three carried a mutation in the FKBP-type peptidyl-prolyl cis-trans isomerase Pfam domain of FPR1, a small peptidylprolyl isomerase that interacts with mTOR. A model of the yeast Tor2/Fpr1/rapamycin tertiary complex shows that residue S1975 is near the bound rapamycin molecule (**Figure 3D**), suggesting that changes at this location might disrupt the formation of the tertiary complex. The model suggests that the two *FPR1* truncation mutations (Y33* and Q61fs) (**Table S4, Figure 3E**) likely confer resistance by interfering with everolimus binding.

**Figure 3:**
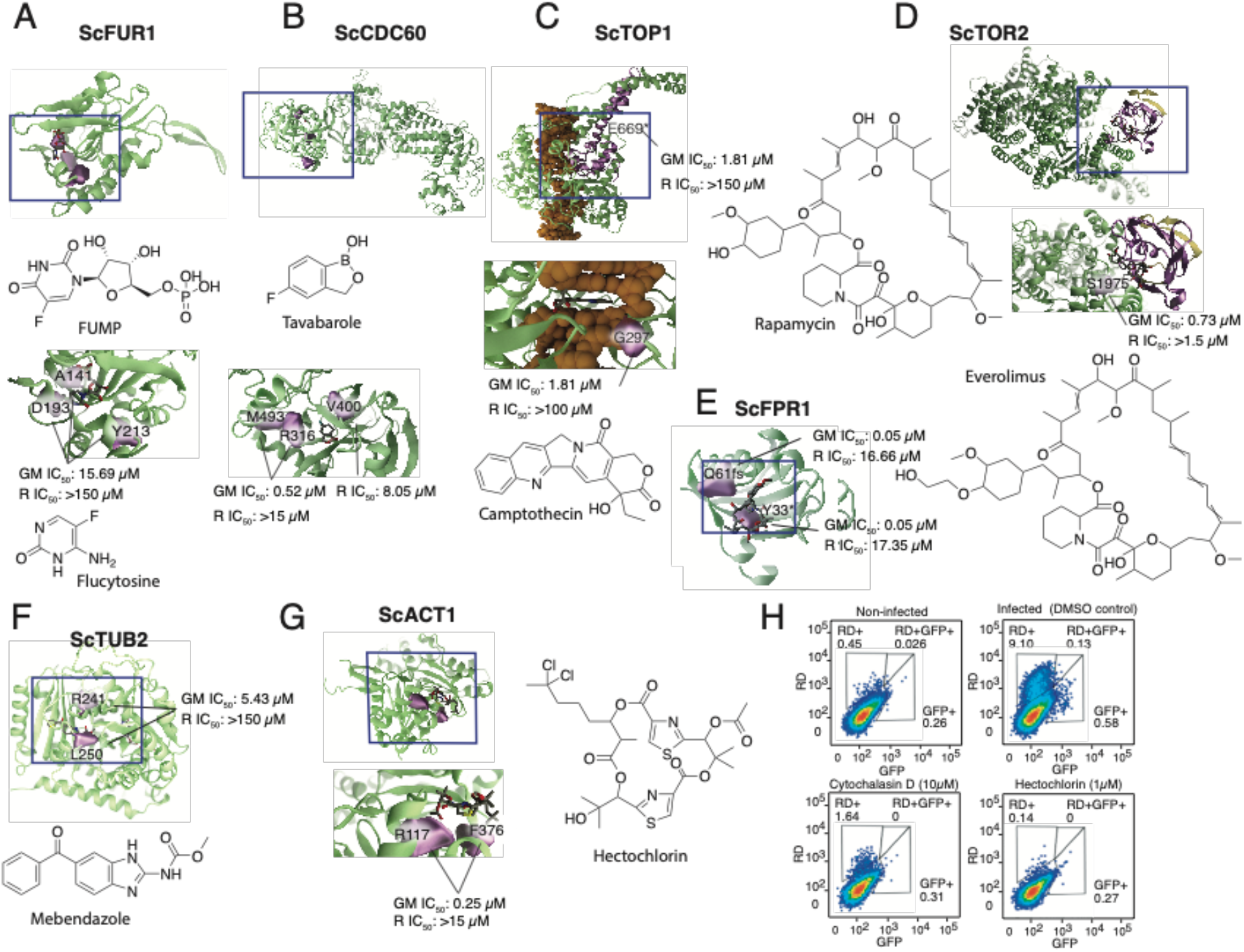
Resistance-conferring mutations in detail. Proteins and DNA are shown in green and orange, respectively. R = resistant line, GM = green monster parents. **A**. **Fur1 in complex with 5FUMP.***Sc*Fur1 homology model, with a bound 5-FUMP analog (uridine monophosphate) taken from an aligned *holo Tm*Fur1 crystal structure (PDB ID: 1O5O). **B**. **Cdc60 model.** Cdc60 homology model bound to a docked tavaborole molecule. **C**. **Top1 model.** DNA-Top1-camptothecin complex, modelled using a *Sc*Top1 crystal structure (PDB: 1OIS), with bound camptothecin taken from an aligned *holo Hs*Top1p crystal structure (PDB: 1T8I). **D**. **Tor2 model.** mTOR-rapamycin-Fpr1 tertiary complex model, modelled using a crystal structure of the human complex as a template (PDB ID: 4DRI^74^). mTOR residues 1001 to 2474 are shown in green (homology model), and Fpr1 is shown in yellow (crystal structure, PDB: 1YAT). **E. Fpr1 model.** Fpr-rapamycin (crystal structure, PDB: 1YAT). **F**. **Tub2 model.** Tub2-nocodazol (crystal structure, PDB: 5CA1). **G**.**Act1 model.** Act1 (crystal structure, PDB: 1YAG), bound to a docked hectochlorin molecule. Evaluation of hepatocellular traversal by P. berghei sporozoites using an established flow cytometry-based assay^75^. **H. Liver cell invasion assay**. Flow cytometry plots show traversal and invasion of host cells at 2 hours post invasion by exoerythrocytic forms in Huh7.5.1 cells. The percent of rhodamine-dextran positive single cells (RD) was used to determine overall traversal frequency, controlled against cytochalasin D (positive) at 10μM and infected untreated conditions, while invasion was evaluated by exclusive GFP+ signal.

Our collection also contained compounds active against other pathogens. For example, mebendazole, a benzimidazole compound, is among the few effective drugs available for treating soil-transmitted helminths (worms) in both humans and animals. It binds to tubulin, thereby disrupting worm motility^28^. We confirmed the antifungal activity^29^ of mebendazole and obtained two resistant strains in our *in vitro* selections. Of the nine missense mutations identified, two were in the GTPase domain of the *TUB2* gene (R241S and L250F) (**Table S4**), near or at the same residues (R241H and R241C) that confer resistance to the related antimitotic drug, benomyl, which also binds tubulin^30,31^. Modeling studies (**Figure 3F**) confirm that the binding mode is similar to that of benomyl, which binds with high affinity to the beta subunit of tubulin, thereby disrupting the structure and function of microtubules^32^. Despite sharing a common target with yeast, helminths and nematodes have benzimidazole-resistance mutations in codons 167, 198 and 200, suggesting some phylum specificity.

Alkylphosphocholines such as miltefosine and edelfosine were originally developed as anticancer agents, but recent work has shown that they are effective against trypanosomatid parasites such as *Leishmania* and *Trypanosoma*^33–36^. The specific target of these drugs remains uncertain. Compound uptake in yeast is known to depend on the membrane transporter Lem3^37,38^, which facilitates phospholipid translocation by interacting with the flippase Dnf1^39^. *DNF1* is closely related to the gene associated with miltefosine resistance in *Leishmania* (Ldmt (AY321297), BLASTP e = 2 × 10^−125^). We identified two independent *LEM3* mutations that confer resistance to miltefosine and edelfosine (K134* and Y107*) (**Table 1**, **Table S4**). Both mutations truncate the protein, functionally mimicking a deletion strain. *LEM3* is a yeast ortholog of LdROS (ABB05176.1, BLASTP 5×10^−13^), also related to *Leishmania* miltefosine resistance.

### Revealing the mechanism of action for uncharacterized compounds

To demonstrate that the yeast model can provide clues about mechanism of action we examined several poorly annotated compounds. Hectochlorin is a natural product from the marine cyanobacterium Lyngbya majuscule^40^ that has strong antimalarial blood stage activity (IC_50_: 85.60 nM ± 0.96) as well as activity against GM yeast (IC_50_=0.25 μM). We identified six independent disruptive mutations, three of which were in the actin Pfam domain of Act1 (*p* = 1.9 ×10^−13^, **Table 1**). We confirmed by CRISPR/Cas9 that a mutation in *ACT1* confers resistance to hectochlorin in yeast (**Table S7**). When mapped onto a crystal structure of Act1 (PDB: 1YAG^41^) the altered amino acids line a distinct protein pocket (**Figure 3G**) suggesting they confer resistance by directly disrupting compound binding. To assess whether hectochlorin resistance in *Plasmodium* occurs through a similar mechanism, we also mapped the mutations onto a synthetic construct of *P. berghei* Act1 protein (PDB: 4CBW (https://www.rcsb.org/structure/4CBW), which shares 97% sequence identity with *Pf*Act1. The altered amino acids again line a well-defined protein pocket, and the hectochlorin docked pose is also similar. This work supports published experiments that suggest actin is the target of hectochlorin^42^. To provide further support for this hypothesis, we determined if hectochlorin produces the same cell-invasion inhibition phenotype in malaria liver stage parasites as cytochalasin D, another actin polymerization inhibitor. Cytochalasin D has been shown to reduce *Plasmodium* sporozoite motility^43^, which is necessary for these exoerythrocytic forms to reach the host liver and begin replication. Treatment with 1 μM hectochlorin effectively blocked parasite invasion as efficiently as 10 μM cytochalasin D (**Figure 3H**).

### Transcriptional mechanisms are associated with multidrug resistance in yeast

Some genes appeared repeatedly across different compound sets. The set of 25 highest confidence genes (mutated five or more times across the dataset) was enriched for DNA-binding transcription factor activity (seven genes, Holm-Bonferonni-corrected *p* = 0.035). Altogether we observed 140 coding mutations in 24 genes affecting transcription (**Figure 4A**). In addition to unique allelic exchanges in *YRR1* (27x) and *YRM1* (23x) (**Figure 4B, C**), multiple unique missense mutations were observed in *PDR1* (7x), *PDR3* (5x) *YAP1* (5x), *AFT1* (5x), *TUP1* (3x) *HAL9* (2x), *AZF1* (2x) (**Table S4)**. With the exception of *TUP1*, *AFT1*, and *YAP1*, all of these genes encode proteins bearing the Zn_2_C_6_ fungal-type DNA-binding domain. In fact, we discovered 124 different mutations in 15 different Zn_2_C_6_ transcription factors. The well-studied transcription factor, Gal4, although not mutated here, is a member of this family. This Zn_2_C_6_ domain family (Pf00172) is only found in fungi: *S. cerevisiae* has 52 genes with this domain, *Candida albicans*, 225. In this human pathogen, members include *FCR1, MRR2, TAC1, PDR1* and *PDR3*, all genes involved in drug resistance. *YRR1 (PDR2), YRM1, PDR1* and *PDR3* are all known to be involved in the pleiotropic drug response in *S. cerevisiae*^44,45^, activating transcription of drug transporters such as *PDR5, PDR10, PDR15, YOR1* and *SNQ2 (reviewed in*^46^). Although the DNA binding is conserved, the central regulatory domains are diverse.

**Figure 4:**
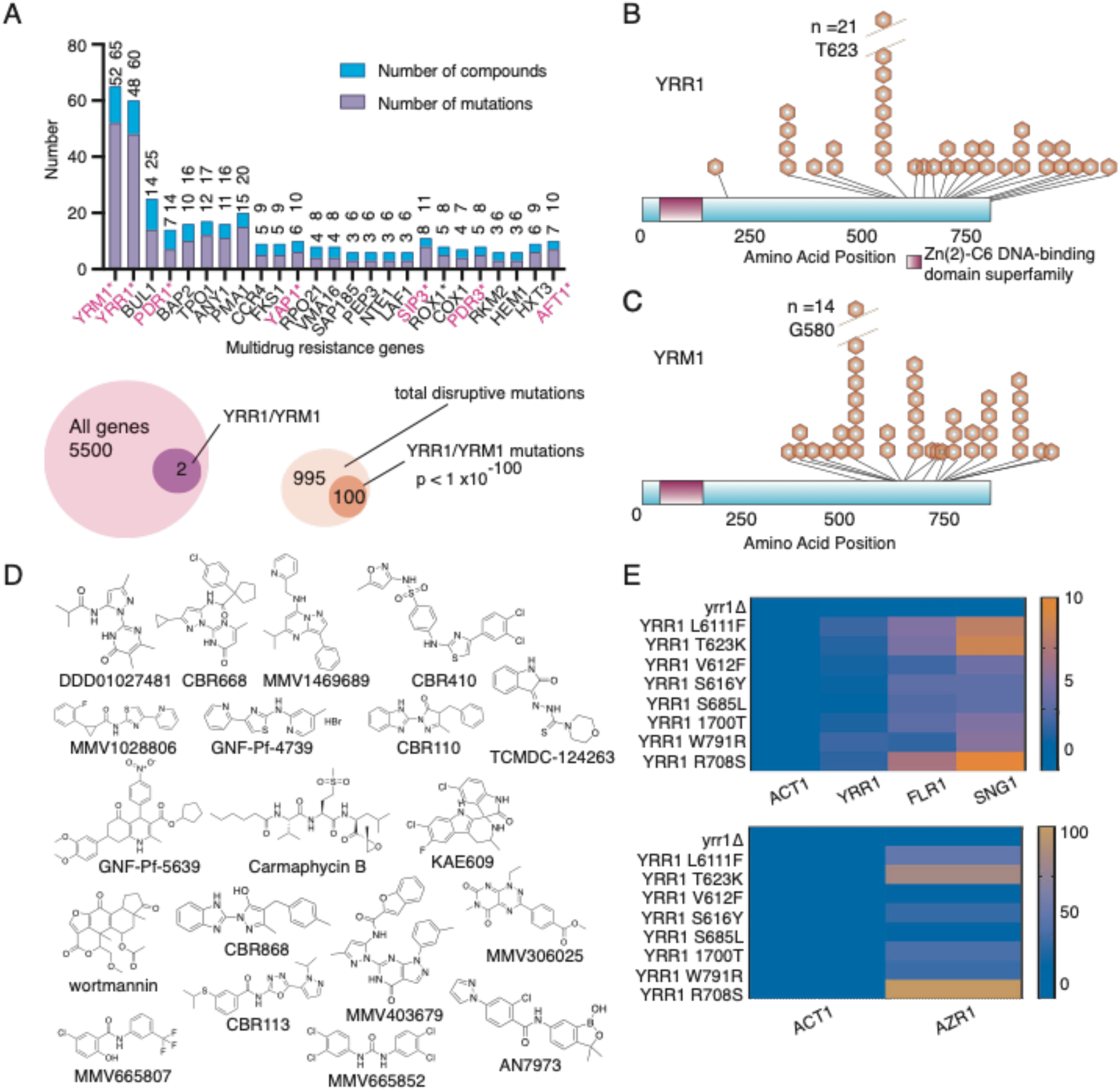
Mutations in transcription factors are over-represented. **A**. **Genes identified with three of more different compounds.** Genes (8) with ontology GO:0140110 (transcription regulator activity) are shown in red. *YRR1* **(B) and***YRM1***(C) mutation localization.** Distribution of mutations in *YRR1* and *YRM1* across the amino acid sequence clustered in the C-terminal activation domain. **D**. **Scaffolds.** Compounds used in selections resulting in *YRR1* and *YRM1* mutations. **E**. *YRR1* **single nucleotide mutations but not loss of function mutations constitutively activate transcriptional targets.** RT-qPCR was utilized to monitor mRNA levels of *YRR1* and *YRR1* activated genes. Strains with *YRR1* point mutations, but not the deletion mutant, show increases in mRNA levels of *YRR1, SNG1, FLG1*, and *AZR1* relative to the wild type strain (GM). In the absence of *YRR1*, associated genes display baseline or lower level of expression. A heatmap legend is shown at right.

The high mutation numbers in *YRR1* and *YRM1 (*which were mutated 100 times for 19, structurally-diverse compounds (**Figure 4B-D**)) allowed us to investigate their spatial localization. Remarkably, all resistance-conferring *YRR1* and *YRM1* mutations were clustered in ~170 amino acids in C terminal half of the protein (**Figure 4B, C**) that is distal to the DNA binding domain. We found no mutations in the DNA binding domain and no mutations in the predicted activation domain at the far C terminal. Notably, it has been shown that the far, C-terminal activation domain of Gal4 (activation domain 9aaTAD (857–871 aa), which interacts with Tra1 protein of the SAGA complex, can be substituted for the activation domain of Yrr1^44^. Yrr1 binds to the sequence (T/A)CCG(C/T)(G/T)(G/T)(A/T)(A/T), found upstream of drug pump genes such as *AZR1, FLR1, SNG1, SNQ2, APD1,* and *PLB1* ^47^.

*YRR1* and *YRM1* are non-essential genes, and we hypothesized that our evolved resistant strains possessed *YRR1* and *YRM1* gain-of-function mutations that result in constitutive expression of transcriptional target genes encoding drug pumps. This hypothesis is motivated in part by the fact that others have reported resistance-conferring gain-of-function mutations in the related genes *PDR1* and *PDR*3^48,49,50^. To expand on this previous work, we exposed a *YRR1* L611F strain (generated using CRISPR/Cas9) to a set of compounds and observed cross-resistance to almost all compounds tested. Of these, compounds that had not previously yielded *YRR1* SNVs in our *in vitro* selections tended to have IC_50_ values only two- to three-fold higher in the *YRR1* L611F strain. In contrast, compounds that had previously yielded *YRR1* mutations in our selections tended to elicit a more than 3-fold IC_50_ difference (**Table S9**).

Others have found that deleting the *YRR1* gene entirely does not significantly increase sensitivity to cytotoxic compounds^48^, providing further evidence that the *YRR1* and *YRM1* mutations identified in our selections—which were all single amino-acid changes—represent gain-of-function mutations. We confirmed this finding by testing a small set of cytotoxic compounds against a *yrr1* deletion strain. With only one exception, we also noted no significant differences in growth inhibition over the wild-type-allele strain. Only compound MMV668507 had a dramatically lower IC_50_ against the *yrr1* deletion strain (**Table S9**).

Based on Gal4 model and previous work^44^ it is likely that the mutated region contains a binding site for one or more repressor proteins and that *YRM1* and *YRR1* SNVs are gain-of-function mutations that result in constitutive expression of drug pump genes. To test this hypothesis, we used qPCR to directly evaluate the expression of three such target genes (*AZR1*, *FLR1,* and *SNG1*). To this set we added the *YRR1* gene itself, since *YRR1* activates its own expression via an auto-feedback loop (**Figure 4E**). We examined gene expression in *YRR1*-mutant strains, in the GM strain with a wild-type *YRR1* allele, and in the *yrr1* deletion strain (**Table S10**). Relative RNA expression levels (**Figure 4E**) of the putative target genes *AZR1*, *FLR1, SNG1* and *YRR1* are 2-70-fold higher in all *YRR1* evolved mutant strains tested compared to the parental GM strain or the *yrr1* deletion strain. To assess whether specific *YRR1* mutations confer resistance to specific compounds or show a more general resistance response, we evaluated nine different *YRR1* mutant strains from our resistance selections (deletion (1), evolved (7), and CRISPR-Cas9 edited (1)) for resistance to a set of four structurally unrelated compounds (CBR113, AN7973, MMV665852, and DDD01027481). All tested strains showed strong cross-resistance to all four compounds (**Table S9**). Taken together, these results strongly support our hypothesis that the identified *YRR1* SNVs lead to constitutive transcriptional activation of their target genes involved in the pleiotropic drug response, thereby conferring general resistance to many compounds. The mutated regions may represent binding sites for repressor proteins that are removed when compounds are present.

### Enrichment of transcription factors is not observed in other species

The repeated identification of transcription factors as mediators of drug resistance in *S. cerevisiae* motivated us to investigate whether similar patterns would be observed in other species. To our knowledge, similar large-scale systematic studies of evolution in the presence of small molecules have only been performed on one other species, the malaria parasite, *Plasmodium falciparum*^51^. In this study, which consisted of evolutions with 37 compounds, mutations were detected in 361 genes. This study showed that many mutations were found directly in a compound’s target and there was no significant enrichment for particular gene class after excluding genes involved in antigenic variation. It is remarkable that in *Plasmodium* only a single nonsynonymous variant in a transcription factor was identified in the entire set of 1905 mutations (although 9 frameshift or inframe indels were observed). Clearly, organisms from the two phyla use different resistance strategies. This may not be unexpected given that *Plasmodium* parasites spend all of their lifecycle within other organisms and are seldom exposed to environmental stresses. These data may suggest why it is relatively easy to kill intracellular parasites with small molecules and why transcriptional profiling may not be useful for drug target identification and mechanism of action studies in some species.

## Discussion

To our knowledge this is one of the most comprehensive studies of the drug selected mutational landscape in fungi. Although genome-wide sets of knockout strains have been used to discover drug targets and to study drug resistance^52,4,53–55^, our approach is different in that we identify single nucleotide gain-of-function mutations in specific domains. This difference allows IVIEWGA to complement other genome-wide knockdown approaches, including those that rely on measures of haploinsufficiency in the presence of compound (HIPHOP)^52^. While *S. cerevisiae* is outstanding model organism, few genetic tools were needed for this study, and the approach can be applied to any organism that can be subjected to drug pressure and sequenced.

One potential disadvantage of whole genome evolutionary approaches is that background passenger mutations can accumulate during the prolonged culturing of a fast-dividing organism and some of these may not contribute to resistance^21^. But given that the overall ratio of nonsynonymous to synonymous changes was 8:1 in our study, most mutations likely do offer some advantage to the cell, even when they are not the primary driver of resistance. For these reasons, a large dataset, such as ours, can provide clarity and statistical confidence, even in the absence of CRISPR-Cas9 reconfirmation of an allele’s importance. The examples that that are discussed in the manuscript did, in almost all cases, achieve strong statistical significance, with mutations appearing in the same genes, or appearing with compounds that are closely related to one another at rates not expected by chance. The reproducibility of the results for genes and compound families indicate that the approach will be powerful in other fungal species that lack the genetic tools that are available for *S. cerevisiae*. It should be mentioned that many, statistically significant examples were not discussed here for the sake of brevity. For example, we find a strong association between *BUL1* and *BAP2* mutations and inhibitors of mitochondrial function and vacuolar ATPases mutations were associated with other scaffold families.

Little was missed in our analysis as well. A resistance allele in one of the 137 genes detected 2 or more times (55 identified more than 2 times) was discovered for 79/80 compounds and in more than 90% of clones. Some of the 30 clones, which did not have a clear, resistance-associated coding SNV had CNVs (e.g. MMV665794-R6a and *ARR1*). In a few cases, they bore mutations in a gene that likely interacts with the gene driving resistance in other clones derived from the same compound treatment: An SNV in *YPK2* was found in one orphan rapamycin-resistant clone (Rap-4R3a) and this is involved in involved in the TORC-dependent phosphorylation of ribosomal proteins. Thus, additional future selections could be useful for defining genetic interaction networks via shared phenotypes.

On the other hand, there may be challenges with extending this approach to other fungal pathogens. Although *S. cerevisiae* is an excellent model it is also haploid. It may be more difficult to apply IVIEWGA to species that are diploid, such as *Candida albicans*. Complete loss of function mutations may be difficult to create and heterozygous gain of function mutations may harder to identify in whole genome sequencing data. Deeper sequencing may be needed for diploids in order to be able reliably call heterozygous alleles. Thus, the mutations that are identified in this study should be useful in interpreting sequencing data from diploid fungal pathogens.

While the enrichment for Zn_2_C_6_ transcription factors in the set of resistance genes may be specific to *Saccharomyces* or even the specific drug pump-depleted GM strain that was used here, smaller-scale studies with specific compounds in fungal pathogens strongly support our findings (reviewed in^56^). Zn_2_C_6_ transcription factors are known to play a major role in the pathogenesis and pleiotropic drug response of pathogenic fungi such as *Candida* spp., the most common clinically relevant fungal pathogens^57,58,59^. Examples include *TAC1*, *STB5* and many others (reviewed in^60^). It is also worthwhile to note that the majority of systematic studies of drug resistance have used yeast knockout collections or gene overexpression systems. Such approaches will favor the identification of drug pumps while missing genes such as *YRR1*, given that knockouts in *YRR1* are not multi-drug resistant.

A major unanswered question is how the cell senses compound increase and translates this to the increases in transcription. It is likely that a cofactor may be binding to the drug resistance domain in Yrr1 and Yrm1, given that these are the location of so many mutations in Zn2C6 transcription factors. Candidates might include Yap1 or Rox1. Yap1 is the most appealing candidate for two reasons—it is one of the few genes repeatedly mutated for compounds that have a general *YRR1/YRM1* response signature but no *YRR1* mutations. It relocalizes to the nucleus in response to cell stress and is involved in multidrug resistance in cancer cells. In addition, others have reported interactions with Yrr1^61^. *YAP1* knockout strains are resistant to a variety of compounds. Yap1 is thought to sense oxidative stress through a cysteine rich domain. Indeed, we found 3 disruptive mutations and 3 missense mutations in *YAP1* and all missense mutations were in cysteines. The ortholog of *YAP1, CAP1*, is also involved in fluconazole resistance in *C. albicans^62^*. Nevertheless, it may be that there are different cofactors for different sets of compounds.

The mutations in *YAP1* highlight how resistance mechanisms are conserved across species and phyla, and while there are clear differences between fungi and other microbes, such as *Plasmodium*: The ortholog of yeast *YAP1*, hsYAP is involved in human cancer drug resistance^63^. Among other genes associated with human chemotherapy resistance include target mutations in TOR2, the target of rapamycin. We show conservation between mutations in *Leishmania* and yeast, *Plasmodium* and yeast as well.

It is noteworthy that none of the clinically approved antifungals (posaconazole, tavaborole, miconazole) are among those that elicited *YRM1-* or *YRR1*-mediated resistance mechanisms, which may indicate why these compounds are ultimately clinically effective against most fungal pathogens. Studies on both the resistance profile and the rate at which resistance emerges are now incorporated into the drug development pipeline for eukaryotic pathogens such as *Plasmodium falciparum* and it may be that studies, similar to those described here, need to be performed if the objective is to create better drugs for fungal pathogens.

## Material and Methods

### Yeast Strains

All yeast strains used are listed in Table S11.

### S. cerevisiae susceptibility and dose-response assays

To measure compound activity against whole-cell yeast, single colonies were inoculated into 2 mL of YPD media and cultured overnight at 250 RPM in a shaking incubator at 30°C. Cultures were diluted the following day and 200 μl of log-phase cultures, (OD_600_nm readings between 0.1 and 0.2) were added to the wells of a 96 well plate. Eight 1:2 serial dilutions were subsequently performed, in biological duplicates, with starting IC_50_ values of 0.15 - 150 μM. After an initial reading of OD_600_ (time 0 hours), the plate was placed in an incubator at 30°C for 18 hours, and OD_600_ nm determined. IC_50_ values were calculated by subtracting OD_600_ nm values at time 0 hours from time 18 hours. As a negative control, cultures not treated with any compounds were run in parallel. Nonlinear regression on log([inhibitor]) vs. response with variable slope was performed using Graphpad Prism.

### Screening the MMV Malaria Box, Pathogen Box, and Charles River libraries

Plates containing 10 μl 10 mM compounds were provided. We first tested these compounds for GM cytotoxicity in single-dose measurements (150 μM, in biological duplicates). Compounds that inhibited GM growth by at least 70% after 18 hours (in either replicate) were further characterized using an eight-point dose-response assay (**Table S1**).

### In vitro resistance evolution

Sublethal compound concentrations were added to *S. cerevisiae* ABC_16_-Monster cells growing logarithmically in YPD media. Each selection culture was grown under vigorous shaking. Upon reaching saturation, cultures were diluted into fresh YPD media containing increasing compound concentrations. Cultures that grew at substantially higher drug concentrations than the parental cell line were streaked for single colonies onto compound-containing agar plates. Single colonies were isolated, and a seven-point dose-response assay (in biological duplicates) with two-fold dilutions was performed to determine the IC_50_ values of the evolved versus parental strains. Genomic DNA from strains that had at least an IC_50_ shift of 1.5-fold was extracted using the YeaStar Genomic DNA kit (cat. No D2002, ZYMO Research).

### Whole-genome sequencing and analysis

Sequencing libraries were prepared using the Illumina Nextera XT kit (Cat. No FC-131-1024, Illumina) following the standard dual indexing protocol, and sequenced on the Illumina HiSeq 2500 in RapidRun mode to generate paired-end reads at least 100 bp in length. Reads were aligned to the *S. cerevisiae* 288C reference genome (assembly R64) using BWA-mem and further processed using Picard Tools (http://broadinstitute.github.io/picard/). Quality control, alignment, and preprocessing workflows were automated using the computational platform Omics Pipe^11^ to ensure scalable and parallelized analysis. A total of 376 clones were sequenced to an average coverage of 55.4x with an average of 99.7% reads mapping to the reference genome. Additional sequencing quality statistics are given in **Table S3**. SNVs and INDELs were called using GATK HaplotypeCaller, filtered based on GATK recommendations^12^, and annotated with SnpEff^13^. Variants were further filtered by removing mutations that were present in both the drug-sensitive parent strain and resistant strains, such that mutations were only retained if they arose during the drug-selection process. Mutations were visually inspected in the Integrative Genomics Viewer (IGV)^64^. Manual annotation of variants was required in some cases to resolve issues with SnpEff outputs. Raw sequencing data files were uploaded to NCBI Sequence Read Archive under accession PRJNA590203. To increase the depth of our analysis we also reanalyzed fastq files from several resistance selections that were previously published (https://escholarship.org/uc/item/42b8231t) and deposited in the NCBI Short Read Archive with the following accession numbers: SRX1745463, SRX1745464, SRX1745465, SRX1745466, SRX1751863, SRX1751950, SRX1751953, SRX1751954, SRX1805319, SRX1805320, SRX1805321, SRX1805322, SRX1805323, SRX1868845, SRX1869272, SRX1869274, SRX1869275, SRX1869276, SRX1869277, SRX1869278, SRX1869279, SRX1869280, SRX1869282). These include selections with the following compounds: KAE609, MMV001239, cycloheximide, MMV000570, MMV007181, MMV019017 and MMV396736.

### CNV Analysis

Coverage values across defined gene intervals in each alignment file were calculated using GATK DiagnoseTargets (input parameters: -max 2000 -ins 1500 -MQ 50). Coverage values were log-transformed then mean-centered across and within arrays in Cluster. Copy number variant were filtered so that they would only be retained if there was at least 2-3x fold coverage change relative to the parent strain and if they spanned four or more genes (**Table S5**). CNVs were visually confirmed in IGV.

### Intergenic mutation analysis

A Python script was written to map the 271 intergenic mutations to known chromosomal features coordinates based on the *S. cerevisiae* S288C genome version R64-2-1 created by the SGD database. Each mutation was located within the chromosomal coordinate of the feature, intron, exon, and other subfeatures, as well as its proximity to those coordinates with a maximum of 500 base pair distant to both up- and downstream directions (**Table S6**).

### CRISPR/Cas9 allelic exchange in S. cerevisiae

CRISPR/Cas9 genome engineering was performed using the *S. cerevisiae* ABC_16_-Monster strain as previously described^65^. gRNA plasmids were generated with specific oligonucleotides (**Table S13**) for the desired allelic exchange (Integrated DNA Technologies) containing a 24 base-pair overlap with the p426 vector backbone. Subsequently, target-specific gRNAs were PCR amplified/transformed into competent *E. coli* cells and selected on LB-Ampicillin plates. ABC_16_-Monster cells expressing Cas9 were simultaneously transformed with 300-500 ng of gene-specific gRNA vector and 1-2 nmole of synthesized donor template (IDT) via a standard lithium acetate method. Transformed cells were plated and selected on methionine and leucine deficient CM-glucose plates. Each engineered mutation was confirmed by Sanger sequencing (Eton Bioscience).

### qPCR

*S. cerevisiae* strains were grown in YPD (1% yeast extract, 2% bacto peptone, 2% dextrose) overnight at 30°C. 1 OD_600_ log-phase cells were harvested and subject to total RNA extraction using Qiagen RNeasy kit, following the manufacturer’s protocol. cDNA was generated using ThermoFisher SuperScript IV First-Strand Synthesis System, following the manufacturer’s protocol using oligo(dT). qPCR was performed with oligonucleotides (**Table S13**) in technical triplicate with Quanta PerfeCTa SYBR® Green FastMix. Analysis was done using Prism 8. Ct values for each gene of interest were averaged and normalized against ACT1 within each strain (ΔCt). Then each gene of interest was normalized against corresponding genes in the wild type GM background (ΔΔCt). Fold expression was calculated using the formula: 2^−ΔΔCt66^. This analysis was done for each of the four biological replicates (**Table S10**).

### Plasmodium invasion assay

The impact of hectochlorin on hepatocellular traversal and invasion by *Plasmodium berghei* (Pb) sporozoites was measured using a previously established flow cytometry-based assay^43^. *Anopheles stephensi* mosquitoes infected with GFP expressing Pb sporozoites (Pb-GFP)^67^ were purchased from the New York University (NYU) Insectary Core Facility. Approximately 24 h before infection, 1.75 × 10^5^ Huh7.5.1 cells were seeded in 24-well plates using DMEM (Invitrogen cat# 11965-092) supplemented with 10% FBS (Corning cat# 35-011-CV) and 1x Pen Strep Glutamine (100 Units/mL Penicillin, 100μg/mL Streptomycin, and 0.292 mg/mL L-glutamine) (Invitrogen cat# 10378-016) for a final volume of 1 mL. On the day of infection, hectochlorin was added to test wells (final concentration 1 μM) with cytochalasin D (final concentration 10 μM) acting as a positive control for invasion inhibition. A non-infected control and DMSO (final concentration 0.5%) negative control was also utilized to mimic the treated well conditions. Sporozoites were freshly dissected and prepared 2-4 h before infection using a previously described method^68^. Immediately prior to infection, rhodamine-dextran was added to each test well (final concentration 1 mg/mL) followed by 3.5 × 10^4^ Pb-GFP sporozoites. The plates were then incubated at 37°C and 5% CO_2_ for 2 h. Following this incubation, the cells were washed and the presence of GFP and rhodamine-dextran signals were evaluated using flow cytometry.

### Model building

We used I-TASSER^25^ to model proteins without acceptable structures in the Protein Data Bank^69^. To visually inspect the homology models, we aligned them to the structural templates used for model construction^70^. We discarded models that had poor I-TASSER C-scores or that we judged to be improbable (e.g., excessively disordered). Where there were no homologous crystal structures with bound ligands for reference, we used docking to predict ligand binding poses. Specifically, we converted the SMILES strings of the ligands to 3D structures using a beta version of Gypsum-DL^71^and docked the 3D models using QuickVina2^23^ (exhaustiveness = 15). The AutoDock forcefield does not include parameters for boron. To dock tavaborole, we substituted the boron atom with a carbon atom, as recommended on the AutoDock webpage (http://autodock.scripps.edu). We tested both the “C” and “A” atom types as boron substitutes to determine which gave predicted tavaborole poses with the best binding affinities. For heme groups, we manually added a charge of +2 to the iron atom. All protein-structure images were generated using BlendMol^72^.

## Supporting information

S4.. Master Mutation Tlist

S5. List of CNVs

S. Library description

S6. List of intergenic mutations

S8. Unconfirmed mutations

S3. Sequencing statistics

S9. YRR1 IC50s

S12. Oligonucleotides

S7. CRISPR IC50s

S2. IC50s of resistant strains

S10. qPCR data

S13. qPCR data

S11. Yeast genotypes

## Acknowledgments

DNA sequencing was performed at the University of California, San Diego (UCSD), with support from the Institute of Genomic Medicine Core. We would like to thank MMV for providing us with the Malaria Box, Pathogen Box, and Charles River Libraries. This study was supported in parts by grants from the Bill and Melinda Gates Foundation (OPP1054480, OPP1141300, OPP1171497) to G.M.G. and E.A.W. and by a National Institutes of Health (NIH) award (GM085764) to E.A.W. and T.I. M.R.L. was supported in part by Ruth L. Kirschstein Institutional National Research Award T32 GM008666 from the National Institute of General Medical Sciences. L.E.C was supported by NIH R01AI120958-01A1. L.E.C. is a Canada Research Chair (Tier 1) in Microbial Genomics & Infectious Disease and co-Director of the CIFAR Fungal Kingdom: Threats & Opportunities program. P.M.K was supported by grant GM127364 from National Institutes of Health. Bill & Melinda Gates Foundation funding (OPP1107194) supported Calibr at Scripps Research for the discovery of the supplied anti-cryptosporidial compounds. This work was supported by a computer-allocation grant from the University of Pittsburgh’s Center for Research Computing to J.D.D.

## Author contributions

E.A.W., S.O., Y.S., P.M.K., L.C., D.F.W., A.K.L., J.D.D., R.M.W. and G.M.G. played roles in conceptualization, project administration, and supervision. J.D.D. and E.H. performed modeling and docking studies, prepared figures and assisted with writing. G.M.G., Y.S., S.O., E.V., P.K., A.L.C., J.S., G.L., J.S. and M.S. performed selections, IC_50_ determinations or CRISPR/Cas9 allelic exchanges. M.R.L., F.G., R.P., R.W. and K.P.G. performed sequence analysis and created figures. K.P.G. performed compound library analysis and created figures and tables. A.K.L., D.F.W., P.M.K., M.T., L.E.C. and L.W. analyzed results. A set of compounds was sourced and provided by W.H.G., M.S.L. and C.W.M. qPCR experiments were performed and analyzed by K.C. Liver cell invasion and FACS assays were done by M.A. T. I. advised on experiments. S.O., E.A.W. and M.R.L. analyzed data, prepared figures and tables and wrote the manuscript. All authors read and approved the submission.

## Data and materials availability

Raw DNA sequences for all 356 yeast strains have been deposited in the Sequence Read Archive (www.ncbi.nlm.nih.gov/sra) under BioProject accession PRJNA590203. All other data are available in the manuscript or the supplementary materials. This work is licensed under a Creative Commons Attribution 4.0 International (CC BY 4.0) license, which permits unrestricted use, distribution, and reproduction in any medium, provided the original work is properly cited. To view a copy of this license, visit http://creativecommons.org/licenses/by/4.0/. This license does not apply to figures/ photos/artwork or other content included in the article that is credited to a third party; obtain authorization from the rights holder before using such material.

## Supplementary Materials

**Table S1. Library description and enriched clusters. Attached Excel spreadsheet.**

**Table S2. IC_50_’s of resistant strains. Attached Excel spreadsheet.**

**Table S3. Sequencing statistics. Attached Excel spreadsheet.**

**Table S4. Master mutation list. Attached Excel spreadsheet.**

**Table S5. List of CNVs. Attached Excel spreadsheet.**

**Table S6. List of intergenic mutations. Attached Excel spreadsheet.**

**Table S7. CRISPR/Cas9 confirmation IC50s. Attached Excel spreadsheet**

**Table S10. qPCR data. Attached Excel spreadsheet**

**Table S12. Oligos used with CRISPR/Cas9 experiments. Attached Excel spreadsheet.**

**Table S8.** Unconfirmed CRISPR/Cas9 mutations.

| GENE | STRAIN NAME | AMINO ACID CHANGE | BP CHANGE | COMPOUND | GM IC<sub>50</sub> ± SD (μM) | CRISPR CLONE IC<sub>50</sub> ± SD (μM) | FC |
| --- | --- | --- | --- | --- | --- | --- | --- |
| **CBR1\*** | EAW521 | G103A | gGa/gCa | lomerizine | 14.71±2.00 | 15.31±8.08 | 1.04 |
| **CDC39** | EAW519 | V825I | Gtt/Att | MMV1469689 | 1.17±1.64 | 0.46±0.43 | 0.39 |
| **HEM1** | EAW274 | Y444S | tAc/tCc | everolimus | 0.05±0.03 | 0.05±0.03 | 1.11 |
| **LSB6** | EAW522 | Q596* | Caa/Taa | sertraline | 14.25±4.69 | 15.38±5.30 | 1.08 |
| **NUP57\*** | EAW490 | A233G | gCg/gGg | CBR668 | 2.29±0.13 | 2.16±0.37 | 0.94 |
| **PMS1** | EAW508 | K724* | Aaa/Taa | doxorubicin | 9.87±1.12 | 7.23±3.09 | 0.73 |
| **POL2** | EAW480 | C1089F | tGt/tTt | lomerizine | 14.71±2.00 | 18.72±12.42 | 1.27 |
| **RPO21** | EAW518 | V462F | Gtt/Ttt | MMV1469689 | 1.17±1.64 | 1.13±1.24 | 0.97 |
| **SGD1** | EAW509 | T370K | aCa/aAa | TCMDC-124263 | 17.54±4.37 | 16.33±4.33 | 0.93 |
| **SIP3** | EAW489 | E704* | Gaa/Taa | GNF-Pf-445 | 9.70±2.64 | 10.95±1.44 | 1.13 |
| **SWH1** | EAW403 | N412K | aaC/aaG | CBR868 | 13.30±3.65 | 6.32±0.76 | 0.47 |
| **TAP42** | EAW287 | T116K | aCa/aAa | MMV403679 | 7.13±1.25 | 3.65±2.02 | 0.51 |
| **TAP42** | EAW287 | T116K | aCa/aAa | CBR668 | 2.29±0.13 | 1.48±0.05 | 0.64 |
| **TCO89** | EAW505 | L293* | tTg/tAg | MMV1469689 | 1.17±1.64 | 0.11±0.05 | 0.09 |
| **TFC1** | EAW275 | Q53K | Cag/Aag | GNF-Pf-2740 | 1.81±0.12 | 1.75±0.38 | 0.97 |
| **TFC4** | EAW280 | D110E | gaC/gaA | GNF-Pf-2740 | 1.81±0.12 | 0.91±0.08 | 0.51 |
| **TSC1** | EAW506 | K100Q | Aag/Cag | TCMDC-124263 | 17.54±4.37 | 20.23±3.40 | 1.15 |
| **UTP18** | EAW363 | G454S | Ggt/Agt | MMV665852 | 0.78±0.34 | 0.20±0.28 | 0.26 |
| **UTP18** | EAW363 | G454S | Ggt/Agt | CBR868 | 13.30±3.65 | 7.10±1.17 | 0.53 |
| **VPS30** | EAW468 | T456K | aCg/aAg | MMV665909 | 8.44±1.50 | 6.64±0.66 | 0.79 |

List of CRISPR/Cas9 mutations that did not confer statistically-significant reduction (less than 1.6x fold-shift) decrease between the edited and parent line. FC, fold change; SD; standard deviation. Data represents the mean of n =3 measurements.

**Table S9.**
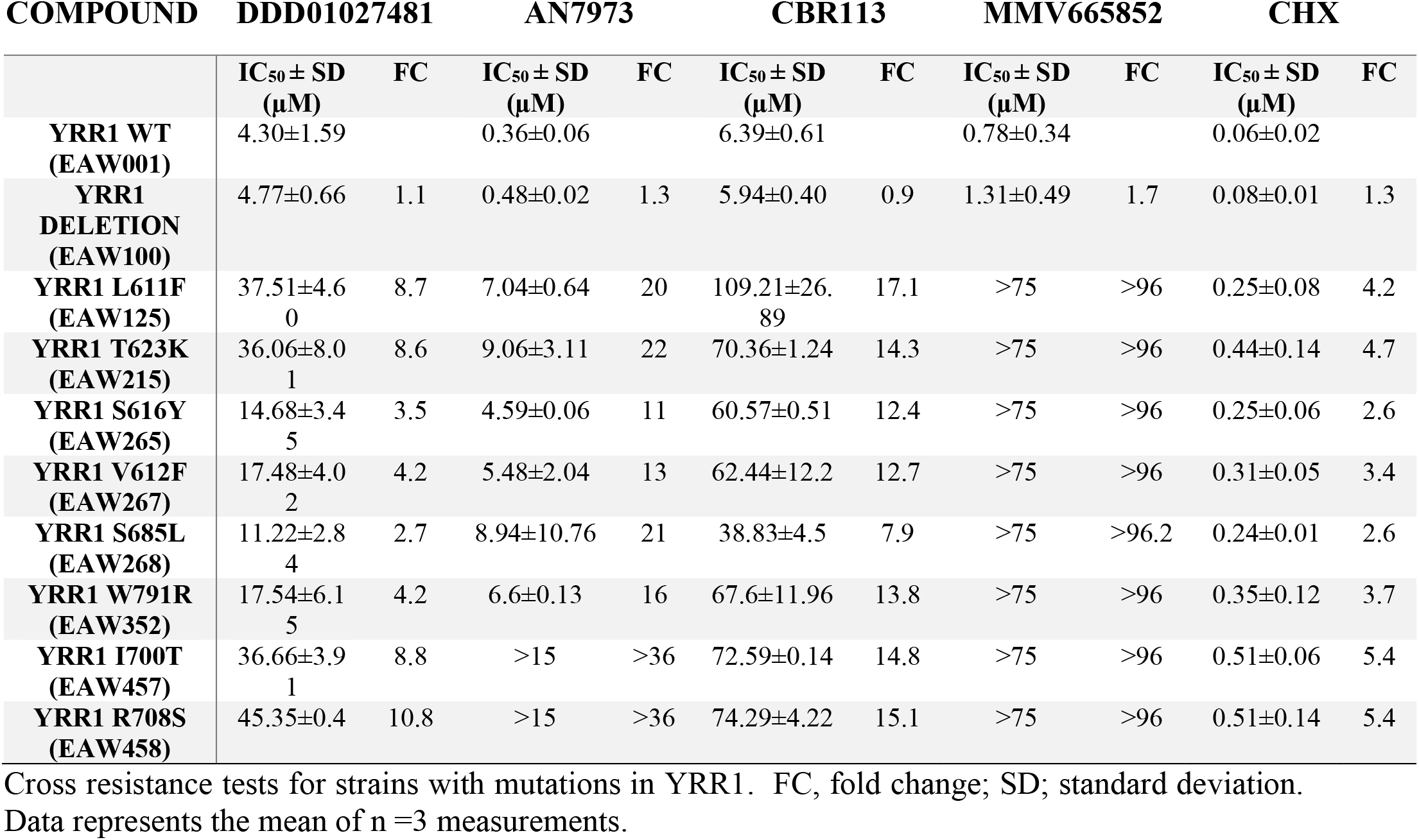
YRR1 strain IC_50_’s.

| COMPOUND | DDD01027481 |  | AN7973 |  | CBR113 |  | MMV665852 |  | CHX |  |
| --- | --- | --- | --- | --- | --- | --- | --- | --- | --- | --- |
|  | IC<sub>50</sub> ± SD<br>(μM) | FC | IC<sub>50</sub> ± SD<br>(μM) | FC | IC<sub>50</sub> ± SD<br>(μM) | FC | IC<sub>50</sub> ± SD<br>(μM) | FC | IC<sub>50</sub> ± SD<br>(μM) | FC |
| **YRR1 WT**<br>**(EAW001)** | 4.30±1.59 |  | 0.36±0.06 |  | 6.39±0.61 |  | 0.78±0.34 |  | 0.06±0.02 |  |
| **YRR1**<br>**DELETION**<br>**(EAW100)** | 4.77±0.66 | 1.1 | 0.48±0.02 | 1.3 | 5.94±0.40 | 0.9 | 1.31±0.49 | 1.7 | 0.08±0.01 | 1.3 |
| **YRR1 L611F**<br>**(EAW125)** | 37.51±4.6<br>0 | 8.7 | 7.04±0.64 | 20 | 109.21±26.<br>89 | 17.1 | >75 | >96 | 0.25±0.08 | 4.2 |
| **YRR1 T623K**<br>**(EAW215)** | 36.06±8.0<br>1 | 8.6 | 9.06±3.11 | 22 | 70.36±1.24 | 14.3 | >75 | >96 | 0.44±0.14 | 4.7 |
| **YRR1 S616Y**<br>**(EAW265)** | 14.68±3.4<br>5 | 3.5 | 4.59±0.06 | 11 | 60.57±0.51 | 12.4 | >75 | >96 | 0.25±0.06 | 2.6 |
| **YRR1 V612F**<br>**(EAW267)** | 17.48±4.0<br>2 | 4.2 | 5.48±2.04 | 13 | 62.44±12.2 | 12.7 | >75 | >96 | 0.31±0.05 | 3.4 |
| **YRR1 S685L**<br>**(EAW268)** | 11.22±2.8<br>4 | 2.7 | 8.94±10.76 | 21 | 38.83±4.5 | 7.9 | >75 | >96.2 | 0.24±0.01 | 2.6 |
| **YRR1 W791R**<br>**(EAW352)** | 17.54±6.1<br>5 | 4.2 | 6.6±0.13 | 16 | 67.6±11.96 | 13.8 | >75 | >96 | 0.35±0.12 | 3.7 |
| **YRR1 I700T**<br>**(EAW457)** | 36.66±3.9<br>1 | 8.8 | >15 | >36 | 72.59±0.14 | 14.8 | >75 | >96 | 0.51±0.06 | 5.4 |
| **YRR1 R708S**<br>**(EAW458)** | 45.35±0.4 | 10.8 | >15 | >36 | 74.29±4.22 | 15.1 | >75 | >96 | 0.51±0.14 | 5.4 |

Cross resistance tests for strains with mutations in YRR1. FC, fold change; SD; standard deviation. Data represents the mean of n =3 measurements.

**Table S11.**
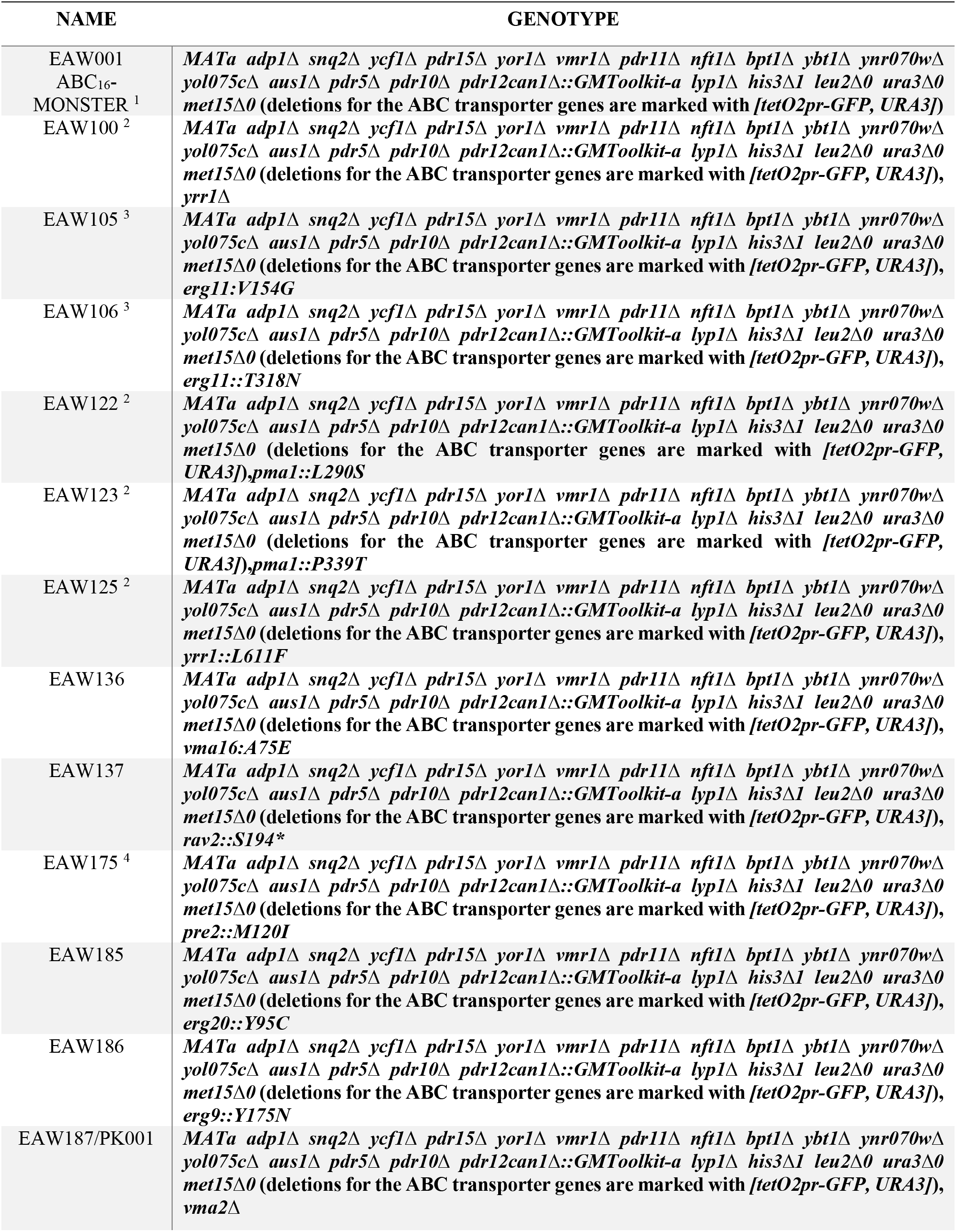
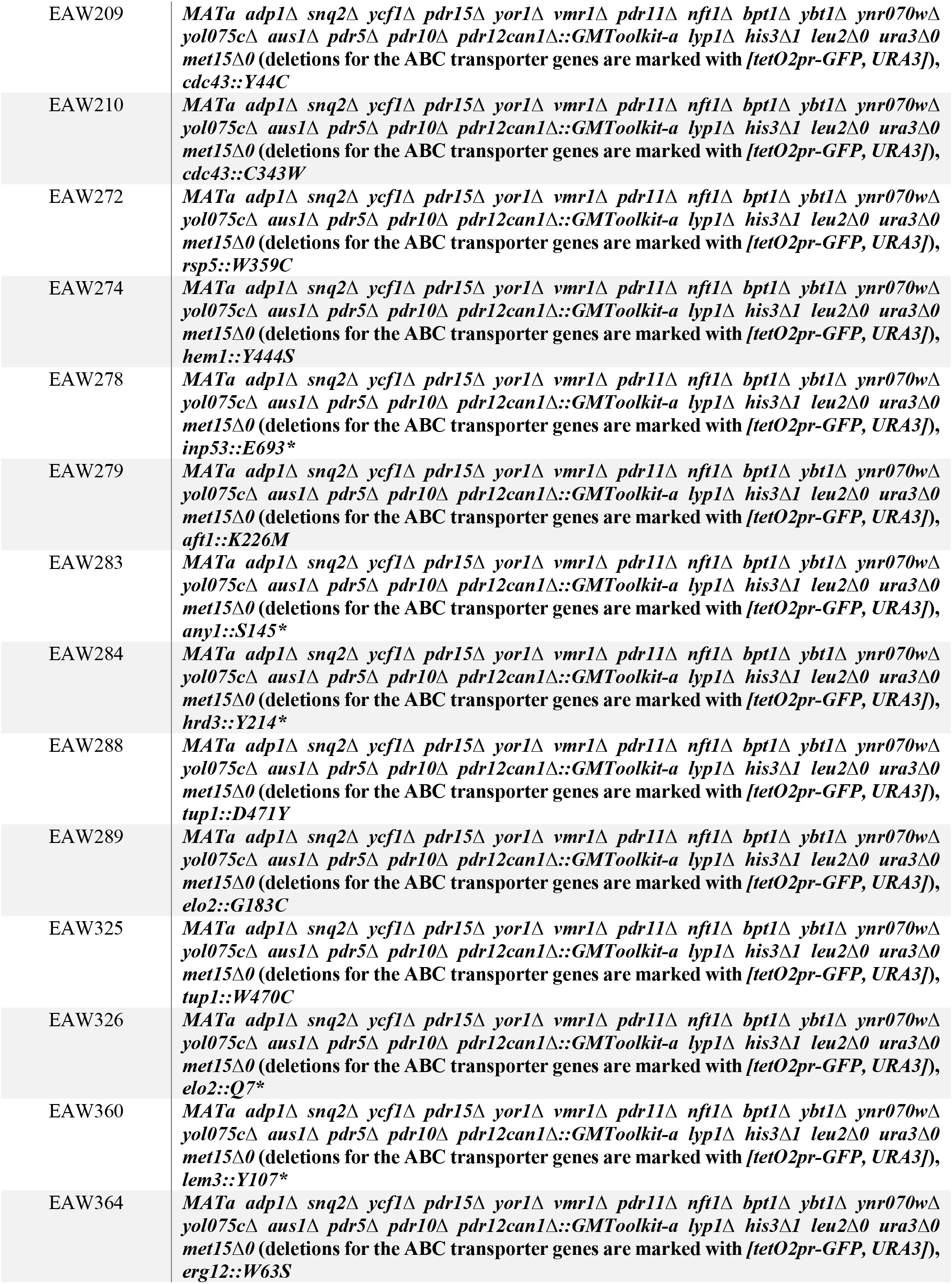
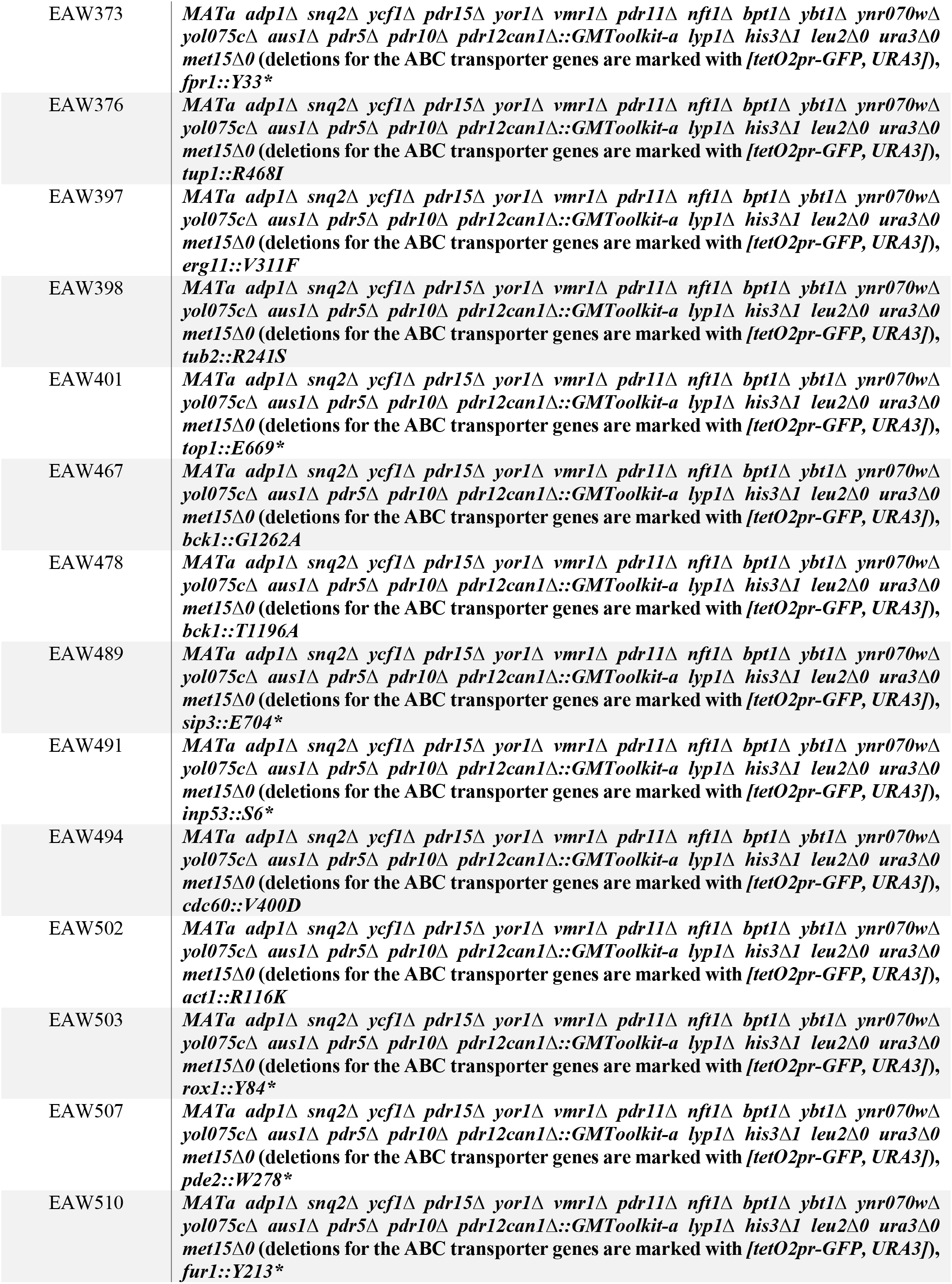
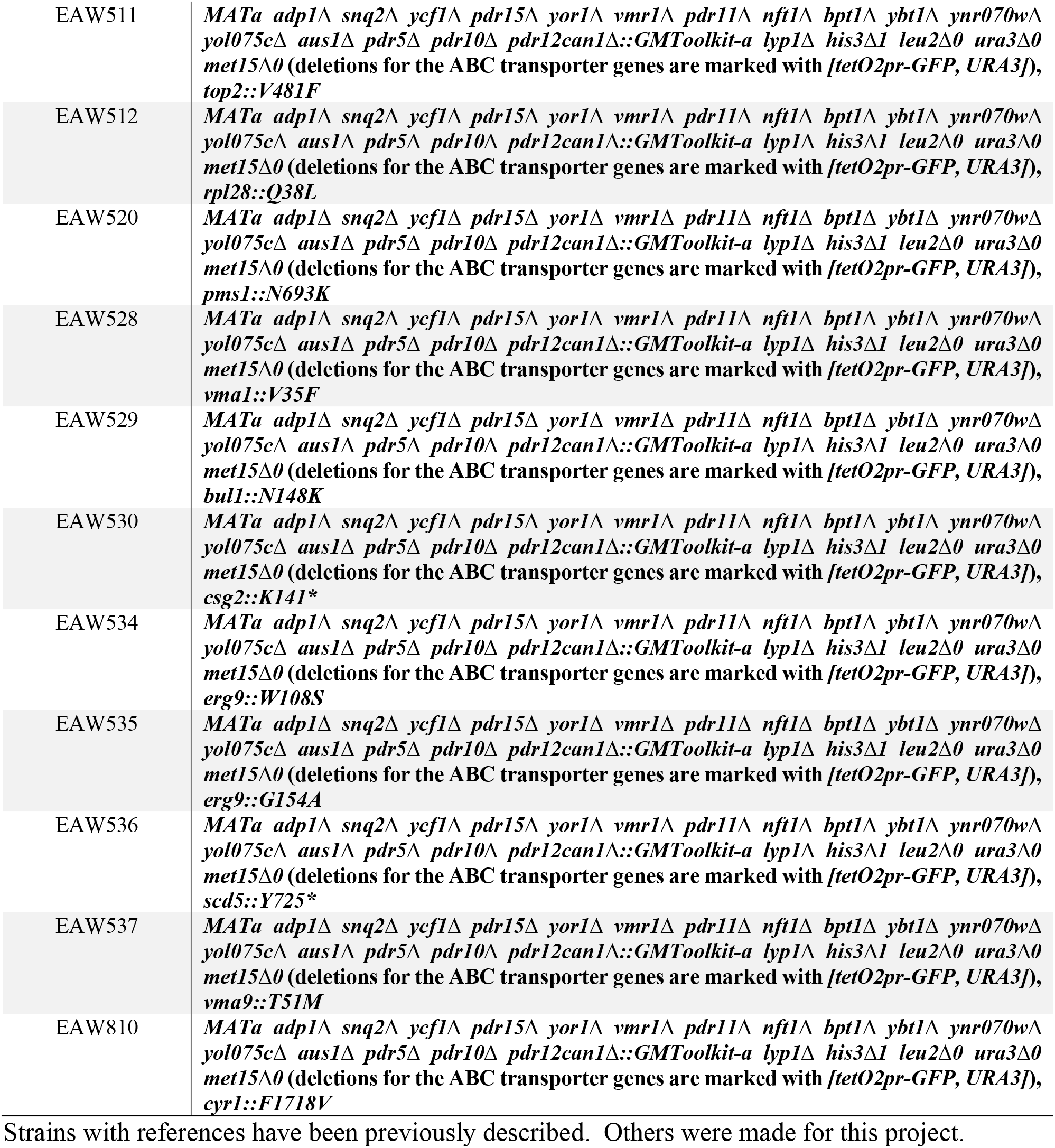
Yeast strain genotype.

**Table S13.**
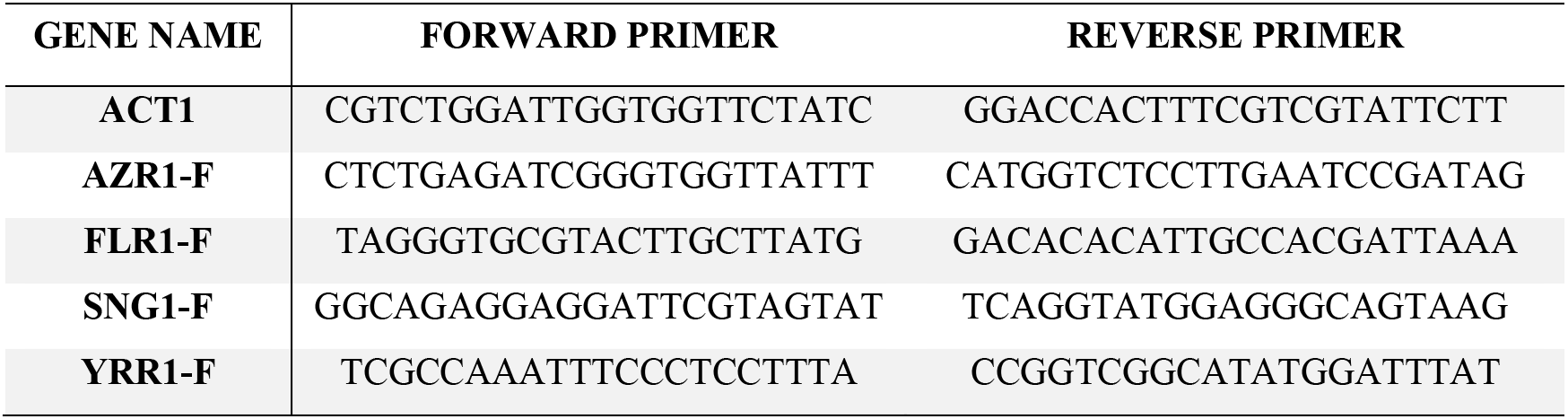
Oligos used with qPCR experiments.

