## Supplementary material for "Defining the Yeast Resistome through *in vitro* Evolution and Whole Genome Sequencing": S11. Yeast genotypes

**Table S11.** Yeast strains used for this study.

| NAME | GENOTYPE |
|---|---|
| EAW001 $ABC_{16}$-MONSTER [1] | ***MATa adp1Δ snq2Δ ycf1Δ pdr15Δ yor1Δ vmr1Δ pdr11Δ nft1Δ bpt1Δ ybt1Δ ynr070wΔ yol075cΔ aus1Δ pdr5Δ pdr10Δ pdr12can1Δ::GMToolkit-a lyp1Δ his3Δ1 leu2Δ0 ura3Δ0 met15Δ0*** **(deletions for the ABC transporter genes are marked with *[tetO2pr-GFP, URA3]*)** |
| EAW100 [2] | ***MATa adp1Δ snq2Δ ycf1Δ pdr15Δ yor1Δ vmr1Δ pdr11Δ nft1Δ bpt1Δ ybt1Δ ynr070wΔ yol075cΔ aus1Δ pdr5Δ pdr10Δ pdr12can1Δ::GMToolkit-a lyp1Δ his3Δ1 leu2Δ0 ura3Δ0 met15Δ0*** **(deletions for the ABC transporter genes are marked with *[tetO2pr-GFP, URA3]*), *yrr1Δ*** |
| EAW105 [3] | ***MATa adp1Δ snq2Δ ycf1Δ pdr15Δ yor1Δ vmr1Δ pdr11Δ nft1Δ bpt1Δ ybt1Δ ynr070wΔ yol075cΔ aus1Δ pdr5Δ pdr10Δ pdr12can1Δ::GMToolkit-a lyp1Δ his3Δ1 leu2Δ0 ura3Δ0 met15Δ0*** **(deletions for the ABC transporter genes are marked with *[tetO2pr-GFP, URA3]*), *erg11:V154G*** |
| EAW106 [3] | ***MATa adp1Δ snq2Δ ycf1Δ pdr15Δ yor1Δ vmr1Δ pdr11Δ nft1Δ bpt1Δ ybt1Δ ynr070wΔ yol075cΔ aus1Δ pdr5Δ pdr10Δ pdr12can1Δ::GMToolkit-a lyp1Δ his3Δ1 leu2Δ0 ura3Δ0 met15Δ0*** **(deletions for the ABC transporter genes are marked with *[tetO2pr-GFP, URA3]*), *erg11::T318N*** |
| EAW122 [2] | ***MATa adp1Δ snq2Δ ycf1Δ pdr15Δ yor1Δ vmr1Δ pdr11Δ nft1Δ bpt1Δ ybt1Δ ynr070wΔ yol075cΔ aus1Δ pdr5Δ pdr10Δ pdr12can1Δ::GMToolkit-a lyp1Δ his3Δ1 leu2Δ0 ura3Δ0 met15Δ0*** **(deletions for the ABC transporter genes are marked with *[tetO2pr-GFP, URA3]*),*pma1::L290S*** |
| EAW123 [2] | ***MATa adp1Δ snq2Δ ycf1Δ pdr15Δ yor1Δ vmr1Δ pdr11Δ nft1Δ bpt1Δ ybt1Δ ynr070wΔ yol075cΔ aus1Δ pdr5Δ pdr10Δ pdr12can1Δ::GMToolkit-a lyp1Δ his3Δ1 leu2Δ0 ura3Δ0 met15Δ0*** **(deletions for the ABC transporter genes are marked with *[tetO2pr-GFP, URA3]*),*pma1::P339T*** |
| EAW125 [2] | ***MATa adp1Δ snq2Δ ycf1Δ pdr15Δ yor1Δ vmr1Δ pdr11Δ nft1Δ bpt1Δ ybt1Δ ynr070wΔ yol075cΔ aus1Δ pdr5Δ pdr10Δ pdr12can1Δ::GMToolkit-a lyp1Δ his3Δ1 leu2Δ0 ura3Δ0 met15Δ0*** **(deletions for the ABC transporter genes are marked with *[tetO2pr-GFP, URA3]*), *yrr1::L611F*** |
| EAW136 | ***MATa adp1Δ snq2Δ ycf1Δ pdr15Δ yor1Δ vmr1Δ pdr11Δ nft1Δ bpt1Δ ybt1Δ ynr070wΔ yol075cΔ aus1Δ pdr5Δ pdr10Δ pdr12can1Δ::GMToolkit-a lyp1Δ his3Δ1 leu2Δ0 ura3Δ0 met15Δ0*** **(deletions for the ABC transporter genes are marked with *[tetO2pr-GFP, URA3]*), *vma16:A75E*** |
| EAW137 | ***MATa adp1Δ snq2Δ ycf1Δ pdr15Δ yor1Δ vmr1Δ pdr11Δ nft1Δ bpt1Δ ybt1Δ ynr070wΔ yol075cΔ aus1Δ pdr5Δ pdr10Δ pdr12can1Δ::GMToolkit-a lyp1Δ his3Δ1 leu2Δ0 ura3Δ0 met15Δ0*** **(deletions for the ABC transporter genes are marked with *[tetO2pr-GFP, URA3]*), *rav2::S194**** |
| EAW175 [4] | ***MATa adp1Δ snq2Δ ycf1Δ pdr15Δ yor1Δ vmr1Δ pdr11Δ nft1Δ bpt1Δ ybt1Δ ynr070wΔ yol075cΔ aus1Δ pdr5Δ pdr10Δ pdr12can1Δ::GMToolkit-a lyp1Δ his3Δ1 leu2Δ0 ura3Δ0 met15Δ0*** **(deletions for the ABC transporter genes are marked with *[tetO2pr-GFP, URA3]*), *pre2::M120I*** |
| EAW185 | ***MATa adp1Δ snq2Δ ycf1Δ pdr15Δ yor1Δ vmr1Δ pdr11Δ nft1Δ bpt1Δ ybt1Δ ynr070wΔ yol075cΔ aus1Δ pdr5Δ pdr10Δ pdr12can1Δ::GMToolkit-a lyp1Δ his3Δ1 leu2Δ0 ura3Δ0 met15Δ0*** **(deletions for the ABC transporter genes are marked with *[tetO2pr-GFP, URA3]*), *erg20::Y95C*** |
| EAW186 | ***MATa adp1Δ snq2Δ ycf1Δ pdr15Δ yor1Δ vmr1Δ pdr11Δ nft1Δ bpt1Δ ybt1Δ ynr070wΔ yol075cΔ aus1Δ pdr5Δ pdr10Δ pdr12can1Δ::GMToolkit-a lyp1Δ his3Δ1 leu2Δ0 ura3Δ0 met15Δ0*** **(deletions for the ABC transporter genes are marked with *[tetO2pr-GFP, URA3]*), *erg9::Y175N*** |
| EAW187/PK001 | ***MATa adp1Δ snq2Δ ycf1Δ pdr15Δ yor1Δ vmr1Δ pdr11Δ nft1Δ bpt1Δ ybt1Δ ynr070wΔ yol075cΔ aus1Δ pdr5Δ pdr10Δ pdr12can1Δ::GMToolkit-a lyp1Δ his3Δ1 leu2Δ0 ura3Δ0*** |

| | |
|---|---|
| | met15Δ0 (deletions for the ABC transporter genes are marked with [tetO2pr-GFP, URA3]), vma2Δ |
| EAW209 | MATa adp1Δ snq2Δ ycf1Δ pdr15Δ yor1Δ vmr1Δ pdr11Δ nft1Δ bpt1Δ ybt1Δ ynr070wΔ yol075cΔ aus1Δ pdr5Δ pdr10Δ pdr12can1Δ::GMToolkit-a lyp1Δ his3Δ1 leu2Δ0 ura3Δ0 met15Δ0 (deletions for the ABC transporter genes are marked with [tetO2pr-GFP, URA3]), cdc43::Y44C |
| EAW210 | MATa adp1Δ snq2Δ ycf1Δ pdr15Δ yor1Δ vmr1Δ pdr11Δ nft1Δ bpt1Δ ybt1Δ ynr070wΔ yol075cΔ aus1Δ pdr5Δ pdr10Δ pdr12can1Δ::GMToolkit-a lyp1Δ his3Δ1 leu2Δ0 ura3Δ0 met15Δ0 (deletions for the ABC transporter genes are marked with [tetO2pr-GFP, URA3]), cdc43::C343W |
| EAW272 | MATa adp1Δ snq2Δ ycf1Δ pdr15Δ yor1Δ vmr1Δ pdr11Δ nft1Δ bpt1Δ ybt1Δ ynr070wΔ yol075cΔ aus1Δ pdr5Δ pdr10Δ pdr12can1Δ::GMToolkit-a lyp1Δ his3Δ1 leu2Δ0 ura3Δ0 met15Δ0 (deletions for the ABC transporter genes are marked with [tetO2pr-GFP, URA3]), rsp5::W359C |
| EAW274 | MATa adp1Δ snq2Δ ycf1Δ pdr15Δ yor1Δ vmr1Δ pdr11Δ nft1Δ bpt1Δ ybt1Δ ynr070wΔ yol075cΔ aus1Δ pdr5Δ pdr10Δ pdr12can1Δ::GMToolkit-a lyp1Δ his3Δ1 leu2Δ0 ura3Δ0 met15Δ0 (deletions for the ABC transporter genes are marked with [tetO2pr-GFP, URA3]), hem1::Y444S |
| EAW278 | MATa adp1Δ snq2Δ ycf1Δ pdr15Δ yor1Δ vmr1Δ pdr11Δ nft1Δ bpt1Δ ybt1Δ ynr070wΔ yol075cΔ aus1Δ pdr5Δ pdr10Δ pdr12can1Δ::GMToolkit-a lyp1Δ his3Δ1 leu2Δ0 ura3Δ0 met15Δ0 (deletions for the ABC transporter genes are marked with [tetO2pr-GFP, URA3]), inp53::E693* |
| EAW279 | MATa adp1Δ snq2Δ ycf1Δ pdr15Δ yor1Δ vmr1Δ pdr11Δ nft1Δ bpt1Δ ybt1Δ ynr070wΔ yol075cΔ aus1Δ pdr5Δ pdr10Δ pdr12can1Δ::GMToolkit-a lyp1Δ his3Δ1 leu2Δ0 ura3Δ0 met15Δ0 (deletions for the ABC transporter genes are marked with [tetO2pr-GFP, URA3]), aft1::K226M |
| EAW283 | MATa adp1Δ snq2Δ ycf1Δ pdr15Δ yor1Δ vmr1Δ pdr11Δ nft1Δ bpt1Δ ybt1Δ ynr070wΔ yol075cΔ aus1Δ pdr5Δ pdr10Δ pdr12can1Δ::GMToolkit-a lyp1Δ his3Δ1 leu2Δ0 ura3Δ0 met15Δ0 (deletions for the ABC transporter genes are marked with [tetO2pr-GFP, URA3]), any1::S145* |
| EAW284 | MATa adp1Δ snq2Δ ycf1Δ pdr15Δ yor1Δ vmr1Δ pdr11Δ nft1Δ bpt1Δ ybt1Δ ynr070wΔ yol075cΔ aus1Δ pdr5Δ pdr10Δ pdr12can1Δ::GMToolkit-a lyp1Δ his3Δ1 leu2Δ0 ura3Δ0 met15Δ0 (deletions for the ABC transporter genes are marked with [tetO2pr-GFP, URA3]), hrd3::Y214* |
| EAW288 | MATa adp1Δ snq2Δ ycf1Δ pdr15Δ yor1Δ vmr1Δ pdr11Δ nft1Δ bpt1Δ ybt1Δ ynr070wΔ yol075cΔ aus1Δ pdr5Δ pdr10Δ pdr12can1Δ::GMToolkit-a lyp1Δ his3Δ1 leu2Δ0 ura3Δ0 met15Δ0 (deletions for the ABC transporter genes are marked with [tetO2pr-GFP, URA3]), tup1::D471Y |
| EAW289 | MATa adp1Δ snq2Δ ycf1Δ pdr15Δ yor1Δ vmr1Δ pdr11Δ nft1Δ bpt1Δ ybt1Δ ynr070wΔ yol075cΔ aus1Δ pdr5Δ pdr10Δ pdr12can1Δ::GMToolkit-a lyp1Δ his3Δ1 leu2Δ0 ura3Δ0 met15Δ0 (deletions for the ABC transporter genes are marked with [tetO2pr-GFP, URA3]), elo2::G183C |
| EAW325 | MATa adp1Δ snq2Δ ycf1Δ pdr15Δ yor1Δ vmr1Δ pdr11Δ nft1Δ bpt1Δ ybt1Δ ynr070wΔ yol075cΔ aus1Δ pdr5Δ pdr10Δ pdr12can1Δ::GMToolkit-a lyp1Δ his3Δ1 leu2Δ0 ura3Δ0 met15Δ0 (deletions for the ABC transporter genes are marked with [tetO2pr-GFP, URA3]), tup1::W470C |
| EAW326 | MATa adp1Δ snq2Δ ycf1Δ pdr15Δ yor1Δ vmr1Δ pdr11Δ nft1Δ bpt1Δ ybt1Δ ynr070wΔ yol075cΔ aus1Δ pdr5Δ pdr10Δ pdr12can1Δ::GMToolkit-a lyp1Δ his3Δ1 leu2Δ0 ura3Δ0 met15Δ0 (deletions for the ABC transporter genes are marked with [tetO2pr-GFP, URA3]), elo2::Q7* |
| EAW360 | MATa adp1Δ snq2Δ ycf1Δ pdr15Δ yor1Δ vmr1Δ pdr11Δ nft1Δ bpt1Δ ybt1Δ ynr070wΔ yol075cΔ aus1Δ pdr5Δ pdr10Δ pdr12can1Δ::GMToolkit-a lyp1Δ his3Δ1 leu2Δ0 ura3Δ0 |

| | |
|---|---|
| | *met15Δ0* (deletions for the ABC transporter genes are marked with *[tetO2pr-GFP, URA3]*), *lem3::Y107** |
| EAW364 | *MATa adp1Δ snq2Δ ycf1Δ pdr15Δ yor1Δ vmr1Δ pdr11Δ nft1Δ bpt1Δ ybt1Δ ynr070wΔ yol075cΔ aus1Δ pdr5Δ pdr10Δ pdr12can1Δ::GMToolkit-a lyp1Δ his3Δ1 leu2Δ0 ura3Δ0 met15Δ0* (deletions for the ABC transporter genes are marked with *[tetO2pr-GFP, URA3]*), *erg12::W63S* |
| EAW373 | *MATa adp1Δ snq2Δ ycf1Δ pdr15Δ yor1Δ vmr1Δ pdr11Δ nft1Δ bpt1Δ ybt1Δ ynr070wΔ yol075cΔ aus1Δ pdr5Δ pdr10Δ pdr12can1Δ::GMToolkit-a lyp1Δ his3Δ1 leu2Δ0 ura3Δ0 met15Δ0* (deletions for the ABC transporter genes are marked with *[tetO2pr-GFP, URA3]*), *fpr1::Y33** |
| EAW376 | *MATa adp1Δ snq2Δ ycf1Δ pdr15Δ yor1Δ vmr1Δ pdr11Δ nft1Δ bpt1Δ ybt1Δ ynr070wΔ yol075cΔ aus1Δ pdr5Δ pdr10Δ pdr12can1Δ::GMToolkit-a lyp1Δ his3Δ1 leu2Δ0 ura3Δ0 met15Δ0* (deletions for the ABC transporter genes are marked with *[tetO2pr-GFP, URA3]*), *tup1::R468I* |
| EAW397 | *MATa adp1Δ snq2Δ ycf1Δ pdr15Δ yor1Δ vmr1Δ pdr11Δ nft1Δ bpt1Δ ybt1Δ ynr070wΔ yol075cΔ aus1Δ pdr5Δ pdr10Δ pdr12can1Δ::GMToolkit-a lyp1Δ his3Δ1 leu2Δ0 ura3Δ0 met15Δ0* (deletions for the ABC transporter genes are marked with *[tetO2pr-GFP, URA3]*), *erg11::V311F* |
| EAW398 | *MATa adp1Δ snq2Δ ycf1Δ pdr15Δ yor1Δ vmr1Δ pdr11Δ nft1Δ bpt1Δ ybt1Δ ynr070wΔ yol075cΔ aus1Δ pdr5Δ pdr10Δ pdr12can1Δ::GMToolkit-a lyp1Δ his3Δ1 leu2Δ0 ura3Δ0 met15Δ0* (deletions for the ABC transporter genes are marked with *[tetO2pr-GFP, URA3]*), *tub2::R241S* |
| EAW401 | *MATa adp1Δ snq2Δ ycf1Δ pdr15Δ yor1Δ vmr1Δ pdr11Δ nft1Δ bpt1Δ ybt1Δ ynr070wΔ yol075cΔ aus1Δ pdr5Δ pdr10Δ pdr12can1Δ::GMToolkit-a lyp1Δ his3Δ1 leu2Δ0 ura3Δ0 met15Δ0* (deletions for the ABC transporter genes are marked with *[tetO2pr-GFP, URA3]*), *top1::E669** |
| EAW467 | *MATa adp1Δ snq2Δ ycf1Δ pdr15Δ yor1Δ vmr1Δ pdr11Δ nft1Δ bpt1Δ ybt1Δ ynr070wΔ yol075cΔ aus1Δ pdr5Δ pdr10Δ pdr12can1Δ::GMToolkit-a lyp1Δ his3Δ1 leu2Δ0 ura3Δ0 met15Δ0* (deletions for the ABC transporter genes are marked with *[tetO2pr-GFP, URA3]*), *bck1::G1262A* |
| EAW478 | *MATa adp1Δ snq2Δ ycf1Δ pdr15Δ yor1Δ vmr1Δ pdr11Δ nft1Δ bpt1Δ ybt1Δ ynr070wΔ yol075cΔ aus1Δ pdr5Δ pdr10Δ pdr12can1Δ::GMToolkit-a lyp1Δ his3Δ1 leu2Δ0 ura3Δ0 met15Δ0* (deletions for the ABC transporter genes are marked with *[tetO2pr-GFP, URA3]*), *bck1::T1196A* |
| EAW489 | *MATa adp1Δ snq2Δ ycf1Δ pdr15Δ yor1Δ vmr1Δ pdr11Δ nft1Δ bpt1Δ ybt1Δ ynr070wΔ yol075cΔ aus1Δ pdr5Δ pdr10Δ pdr12can1Δ::GMToolkit-a lyp1Δ his3Δ1 leu2Δ0 ura3Δ0 met15Δ0* (deletions for the ABC transporter genes are marked with *[tetO2pr-GFP, URA3]*), *sip3::E704** |
| EAW491 | *MATa adp1Δ snq2Δ ycf1Δ pdr15Δ yor1Δ vmr1Δ pdr11Δ nft1Δ bpt1Δ ybt1Δ ynr070wΔ yol075cΔ aus1Δ pdr5Δ pdr10Δ pdr12can1Δ::GMToolkit-a lyp1Δ his3Δ1 leu2Δ0 ura3Δ0 met15Δ0* (deletions for the ABC transporter genes are marked with *[tetO2pr-GFP, URA3]*), *inp53::S6** |
| EAW494 | *MATa adp1Δ snq2Δ ycf1Δ pdr15Δ yor1Δ vmr1Δ pdr11Δ nft1Δ bpt1Δ ybt1Δ ynr070wΔ yol075cΔ aus1Δ pdr5Δ pdr10Δ pdr12can1Δ::GMToolkit-a lyp1Δ his3Δ1 leu2Δ0 ura3Δ0 met15Δ0* (deletions for the ABC transporter genes are marked with *[tetO2pr-GFP, URA3]*), *cdc60::V400D* |
| EAW502 | *MATa adp1Δ snq2Δ ycf1Δ pdr15Δ yor1Δ vmr1Δ pdr11Δ nft1Δ bpt1Δ ybt1Δ ynr070wΔ yol075cΔ aus1Δ pdr5Δ pdr10Δ pdr12can1Δ::GMToolkit-a lyp1Δ his3Δ1 leu2Δ0 ura3Δ0 met15Δ0* (deletions for the ABC transporter genes are marked with *[tetO2pr-GFP, URA3]*), *act1::R116K* |
| EAW503 | *MATa adp1Δ snq2Δ ycf1Δ pdr15Δ yor1Δ vmr1Δ pdr11Δ nft1Δ bpt1Δ ybt1Δ ynr070wΔ yol075cΔ aus1Δ pdr5Δ pdr10Δ pdr12can1Δ::GMToolkit-a lyp1Δ his3Δ1 leu2Δ0 ura3Δ0* |

| | |
|---|---|
| | ***met15Δ0*** **(deletions for the ABC transporter genes are marked with** ***[tetO2pr-GFP, URA3]*****),** ***rox1::Y84**** |
| EAW507 | ***MATa adp1Δ snq2Δ ycf1Δ pdr15Δ yor1Δ vmr1Δ pdr11Δ nft1Δ bpt1Δ ybt1Δ ynr070wΔ yol075cΔ aus1Δ pdr5Δ pdr10Δ pdr12can1Δ::GMToolkit-a lyp1Δ his3Δ1 leu2Δ0 ura3Δ0 met15Δ0*** **(deletions for the ABC transporter genes are marked with** ***[tetO2pr-GFP, URA3]*****),** ***pde2::W278**** |
| EAW510 | ***MATa adp1Δ snq2Δ ycf1Δ pdr15Δ yor1Δ vmr1Δ pdr11Δ nft1Δ bpt1Δ ybt1Δ ynr070wΔ yol075cΔ aus1Δ pdr5Δ pdr10Δ pdr12can1Δ::GMToolkit-a lyp1Δ his3Δ1 leu2Δ0 ura3Δ0 met15Δ0*** **(deletions for the ABC transporter genes are marked with** ***[tetO2pr-GFP, URA3]*****),** ***fur1::Y213**** |
| EAW511 | ***MATa adp1Δ snq2Δ ycf1Δ pdr15Δ yor1Δ vmr1Δ pdr11Δ nft1Δ bpt1Δ ybt1Δ ynr070wΔ yol075cΔ aus1Δ pdr5Δ pdr10Δ pdr12can1Δ::GMToolkit-a lyp1Δ his3Δ1 leu2Δ0 ura3Δ0 met15Δ0*** **(deletions for the ABC transporter genes are marked with** ***[tetO2pr-GFP, URA3]*****),** ***top2::V481F*** |
| EAW512 | ***MATa adp1Δ snq2Δ ycf1Δ pdr15Δ yor1Δ vmr1Δ pdr11Δ nft1Δ bpt1Δ ybt1Δ ynr070wΔ yol075cΔ aus1Δ pdr5Δ pdr10Δ pdr12can1Δ::GMToolkit-a lyp1Δ his3Δ1 leu2Δ0 ura3Δ0 met15Δ0*** **(deletions for the ABC transporter genes are marked with** ***[tetO2pr-GFP, URA3]*****),** ***rpl28::Q38L*** |
| EAW520 | ***MATa adp1Δ snq2Δ ycf1Δ pdr15Δ yor1Δ vmr1Δ pdr11Δ nft1Δ bpt1Δ ybt1Δ ynr070wΔ yol075cΔ aus1Δ pdr5Δ pdr10Δ pdr12can1Δ::GMToolkit-a lyp1Δ his3Δ1 leu2Δ0 ura3Δ0 met15Δ0*** **(deletions for the ABC transporter genes are marked with** ***[tetO2pr-GFP, URA3]*****),** ***pms1::N693K*** |
| EAW528 | ***MATa adp1Δ snq2Δ ycf1Δ pdr15Δ yor1Δ vmr1Δ pdr11Δ nft1Δ bpt1Δ ybt1Δ ynr070wΔ yol075cΔ aus1Δ pdr5Δ pdr10Δ pdr12can1Δ::GMToolkit-a lyp1Δ his3Δ1 leu2Δ0 ura3Δ0 met15Δ0*** **(deletions for the ABC transporter genes are marked with** ***[tetO2pr-GFP, URA3]*****),** ***vma1::V35F*** |
| EAW529 | ***MATa adp1Δ snq2Δ ycf1Δ pdr15Δ yor1Δ vmr1Δ pdr11Δ nft1Δ bpt1Δ ybt1Δ ynr070wΔ yol075cΔ aus1Δ pdr5Δ pdr10Δ pdr12can1Δ::GMToolkit-a lyp1Δ his3Δ1 leu2Δ0 ura3Δ0 met15Δ0*** **(deletions for the ABC transporter genes are marked with** ***[tetO2pr-GFP, URA3]*****),** ***bul1::N148K*** |
| EAW530 | ***MATa adp1Δ snq2Δ ycf1Δ pdr15Δ yor1Δ vmr1Δ pdr11Δ nft1Δ bpt1Δ ybt1Δ ynr070wΔ yol075cΔ aus1Δ pdr5Δ pdr10Δ pdr12can1Δ::GMToolkit-a lyp1Δ his3Δ1 leu2Δ0 ura3Δ0 met15Δ0*** **(deletions for the ABC transporter genes are marked with** ***[tetO2pr-GFP, URA3]*****),** ***csg2::K141**** |
| EAW534 | ***MATa adp1Δ snq2Δ ycf1Δ pdr15Δ yor1Δ vmr1Δ pdr11Δ nft1Δ bpt1Δ ybt1Δ ynr070wΔ yol075cΔ aus1Δ pdr5Δ pdr10Δ pdr12can1Δ::GMToolkit-a lyp1Δ his3Δ1 leu2Δ0 ura3Δ0 met15Δ0*** **(deletions for the ABC transporter genes are marked with** ***[tetO2pr-GFP, URA3]*****),** ***erg9::W108S*** |
| EAW535 | ***MATa adp1Δ snq2Δ ycf1Δ pdr15Δ yor1Δ vmr1Δ pdr11Δ nft1Δ bpt1Δ ybt1Δ ynr070wΔ yol075cΔ aus1Δ pdr5Δ pdr10Δ pdr12can1Δ::GMToolkit-a lyp1Δ his3Δ1 leu2Δ0 ura3Δ0 met15Δ0*** **(deletions for the ABC transporter genes are marked with** ***[tetO2pr-GFP, URA3]*****),** ***erg9::G154A*** |
| EAW536 | ***MATa adp1Δ snq2Δ ycf1Δ pdr15Δ yor1Δ vmr1Δ pdr11Δ nft1Δ bpt1Δ ybt1Δ ynr070wΔ yol075cΔ aus1Δ pdr5Δ pdr10Δ pdr12can1Δ::GMToolkit-a lyp1Δ his3Δ1 leu2Δ0 ura3Δ0 met15Δ0*** **(deletions for the ABC transporter genes are marked with** ***[tetO2pr-GFP, URA3]*****),** ***scd5::Y725**** |
| EAW537 | ***MATa adp1Δ snq2Δ ycf1Δ pdr15Δ yor1Δ vmr1Δ pdr11Δ nft1Δ bpt1Δ ybt1Δ ynr070wΔ yol075cΔ aus1Δ pdr5Δ pdr10Δ pdr12can1Δ::GMToolkit-a lyp1Δ his3Δ1 leu2Δ0 ura3Δ0 met15Δ0*** **(deletions for the ABC transporter genes are marked with** ***[tetO2pr-GFP, URA3]*****),** ***vma9::T51M*** |
| EAW810 | ***MATa adp1Δ snq2Δ ycf1Δ pdr15Δ yor1Δ vmr1Δ pdr11Δ nft1Δ bpt1Δ ybt1Δ ynr070wΔ yol075cΔ aus1Δ pdr5Δ pdr10Δ pdr12can1Δ::GMToolkit-a lyp1Δ his3Δ1 leu2Δ0 ura3Δ0*** |

| | |
|---|---|
| | ***met15Δ0*** **(deletions for the ABC transporter genes are marked with *[tetO2pr-GFP, URA3]*), *cyr1::F1718V*** |
